# The impact of extracellular matrix on the precision medicine utility of pancreatic cancer patient-derived organoids

**DOI:** 10.1101/2023.01.26.525757

**Authors:** Jan C. Lumibao, Shira R. Okhovat, Kristina L. Peck, Xiaoxue Lin, Kathryn Lande, Jingjing Zou, Dannielle D. Engle

**Affiliations:** Salk Institute for Biological Studies; University of California San Diego

**Keywords:** organoids, PDAC, pharmacotyping, precision medicine, basement membrane extract

## Abstract

The use of patient-derived organoids (PDOs) to characterize therapeutic sensitivity and resistance (pharmacotyping) is a promising precision medicine approach. The potential of this approach to inform clinical decisions is now being tested in several large multi-institutional clinical trials. PDOs are cultivated in extracellular matrix from basement membrane extracts (BMEs) that are most commonly acquired commercially. Each clinical site utilizes distinct BME lots and may be restricted due to the availability of commercial BME sources. However, the impact of different sources and lots of BMEs on organoid drug response is unknown. Here, we tested the impact of BME source and lot on proliferation, chemotherapy and targeted therapy drug response, and gene expression in mouse and human pancreatic ductal adenocarcinoma (PDA) organoids. Both human and mouse organoids displayed increased proliferation in Matrigel (Corning) compared to Cultrex (RnD) and UltiMatrix (RnD). However, we observed no substantial impact on drug response when oragnoids were cultured in Matrigel, Cultrex, or UltiMatrix. We also did not observe major shifts in gene expression across the different BME sources, and PDOs maintained their Classical or Basal-like designation. Overall, we find that BME source (Matrigel, Cultrex, UltiMatrix) does not shift PDO dose-response curves and drug testing results, indicating that PDO pharmacotyping is a robust approach for precision medicine.

## Introduction

The 5-year relative survival rate for patients with pancreatic ductal adenocarcinoma (PDA) is 12% (*1*). As the incidence of PDA continues to rise (*2*), strategies to increase survival are imperative. Surgical resection remains the only curative option. However, most patients are ineligible due to the presence of metastasis at the time of diagnosis (*3*). Standard-of-care chemotherapeutic regimens include gemcitabine/nab-paclitaxel and FOLFIRINOX, which demonstrate median survival times of 8.5 and 11.1 months, respectively (*4, 5*). Further, as heterogeneity in tumor biology and treatment response continues to be revealed, personalized medicine approaches are becoming increasingly necessary.

The use of patient-derived organoids (PDOs) has the potential to revolutionize care for patients with PDA (*6*). PDOs are 3D cultures in defined conditions that support propagation of normal, premalignant, and neoplastic cells from primary tissue (*7–11*). Organoid technology has emerged as a promising avenue for precision medicine (*12*). The ability to derive PDA PDOs from surgical resections, rapid autopsies (RAP), and endoscopic ultrasound guided fine needle biopsies allows for a broad sampling of patients with PDA to encompass inter- and intra-patient tumor heterogeneity (*8*). Importantly, PDA PDOs mirror patient tumor genetics, gene expression, and treatment response, situating them as a promising tool for more precision medicine efforts to identify alternative treatment strategies (*8, 13–15*).

As the use of organoids has expanded and become more accessible, variations in culture conditions have been introduced to optimize PDA PDO generation and growth. A number of studies have delineated the impact of the liquid media composition on organoid phenotype, transcriptome, and drug response (*16, 17*). Commercial products to support organoid studies have become more widespread, including a wide variety of basement membrane extracts (BME), which serve as 3D scaffolds. Recent studies have described the use of fully synthetic and defined hydrogels to support organoids of colon, bile duct, and mammary gland origin (*18–21*). However, BMEs derived from Engelbreth-Holm-Swarm (EHS) murine sarcomas, most notably Matrigel (Corning), but also products such as Cultrex and UltiMatrix (RnD), remain the most widely used (*19*). EHS-tumor-derived BMEs contain heterogenous mixtures of extracellular matrix proteins, primarily laminins, collagen IV, entactin, and perlecan, as well as tumor-derived proteins and growth factors, which contribute to batch-to-batch variability (*18, 19*). Regardless, these BMEs have historically served as the scaffold for organoid culture. However, the impact of different EHS- tumor-derived BMEs on organoid growth, chemosensitivity, and global gene expression remains unclear. Importantly, previous reports have demonstrated the clinical utility and relevance of PDA PDO pharmacotyping using Matrigel (*8, 13, 15*). These studies and others have prompted the design of clinical trials based on the use of organoid-guided chemotherapy. However, due to the CoVID-19 pandemic, severe supply chain issues have limited BME availability. These issues range from complete inability to secure the same commercial source of BME or dramatic reduction of available BME within the same lot. The impact of the commercial source and lot of BME on drug response and prognostic gene expression programs has not been explored, yet is critical for the deployment of PDO clinical trials.

Here, we report the impact of BME on PDA organoid growth, response to standard-of-care chemotherapy as well as targeted therapy, and gene expression patterns. While we find that BME source has a significant and substantial impact on organoid growth, drug response and gene expression remain consistent across multiple lots and commercial sources. Results from this study provide insights into selecting BME for organoid culture and for characterizing PDO sensitivity and resistance to both chemotherapies and targeted therapies, which may assist in guiding clinical decisions.

## Results

### BME impacts mouse and human organoid growth

To investigate the influence of BME on organoid growth, human and mouse PDA organoids (**Supplementary Table 1-2**) were cultivated in Matrigel 04 (lot #1062004) and then plated in either Matrigel 04, Matrigel 01 (lot #0287001), Cultrex 83 (lot #1564183), Cultrex 87 (lot #1586187), or UltiMatrix 96 (lot #1637796). Intracellular ATP levels were measured after 4 (mouse) or 6 days of culture (human). Human PDOs and mouse organoids exhibited reduced growth in Cultrex and UltiMatrix compared to Matrigel (**Figure 1A-B, Supplementary Figure 2**). Mouse organoid proliferation did not recover even after multiple passages in Cultrex, suggesting that this growth deficit is not due to a lack of adaptation period (data not shown). In most experimental replicates across mouse and human organoids, there was no difference in proliferation between Matrigel lots. In 1 out of 3 experiments, there was a small but significant decrease (<10%) in growth of hM1F and hF24 in Matrigel 01 compared to Matrigel 04 (**Supplementary Figure 2A-B**). Similarly, 1 out of 2 experiments demonstrated a significant decrease (24%) in hT1 growth in Matrigel 01 (**Supplementary Figure 2C**). Mouse organoid mT69B consistently exhibited no significant differences in intracellular ATP levels between Matrigel lots (n=4/4), while mT69A exhibited 17-23% changes between Matrigel lots (n=2/4) (**Supplementary Figure 2D-E**). In nearly all hPDO experiments, there was a significant, and sometimes substantial (e.g., 27% in Cultrex 83 relative to Matrigel 04), reduction in growth when cultured in either lot of Cultrex or UltiMatrix compared to Matrigel 04 (**Figure 1A, Supplementary Figure 2A-B**). Mouse organoids exhibited more mixed results across experimental repeats, but consistently (n=3/4 mT69A, n=4/4 mT69B) exhibited significant, and sometimes substantial (e.g., 55% decrease in Ultimatrix 96 relative to Matrigel 04), decreased growth in UltiMatrix 96 (**Figure 1B, Supplementary Figure 2D-E**). In the setting of Cultrex, mouse organoid proliferation trended slower relative to Matrigel. mT69A and B grew significantly slower in Cultrex 83 (A: n=2/4, B: 2/4) and 87 (A: n=3/4, B: 2/4), while exhibiting no significant difference relative to Matrigel 04 in 25- 50% of the experiments. These data collectively indicate that mouse and human organoids grow significantly more in Matrigel compared to Cultrex or UltiMatrix. The impact of BME source on PDO expansion may be potentially problematic for completion of drug testing for patients within a clinically relevant timeline.

**Figure 1.**
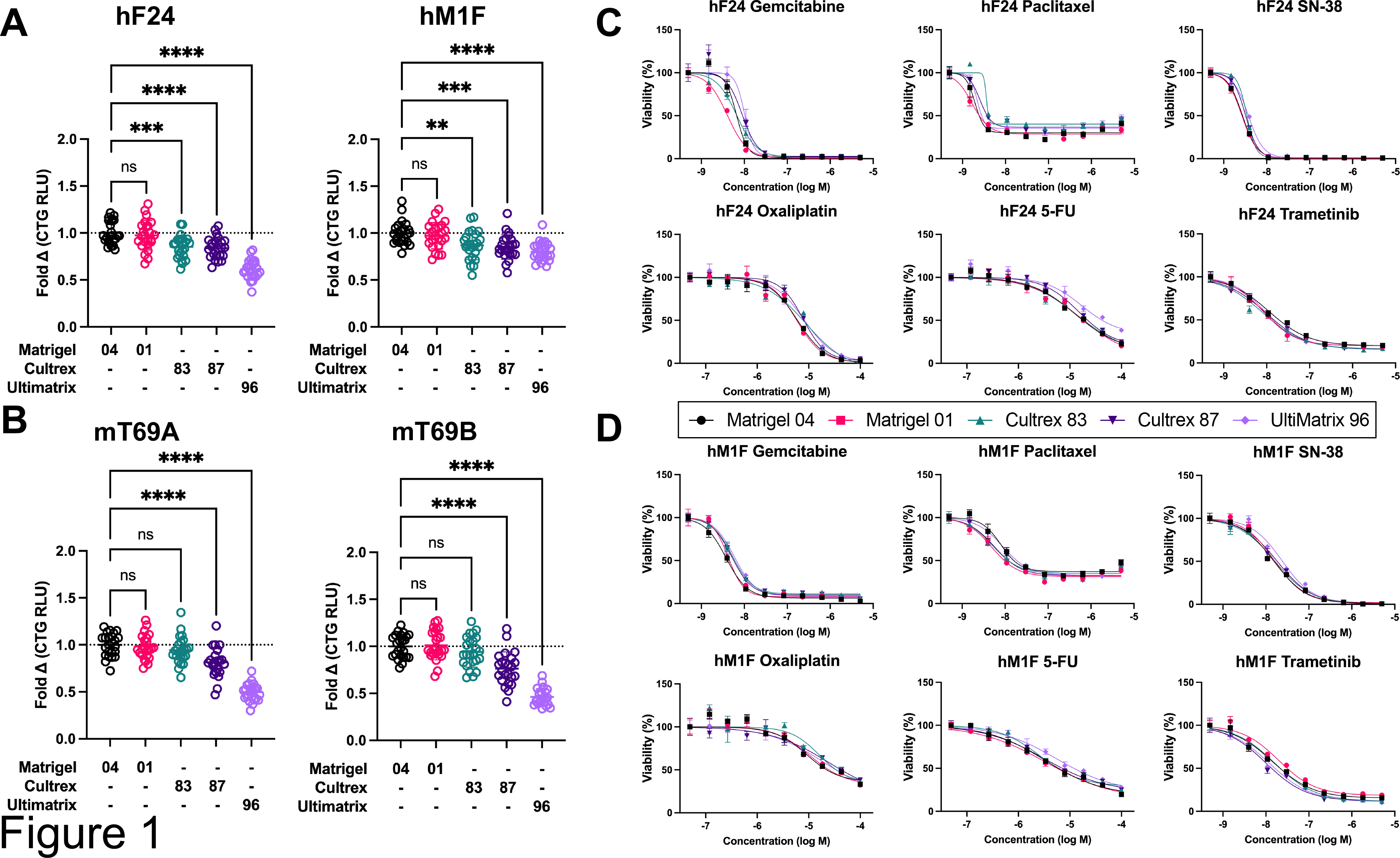
Basement membrane extract impacts growth of mouse and human organoids but does not significantly shift dose-response curves. (A) Growth of human patient-derived organoids hF24 and hM1F cultured in Matrigel 04, Matrigel 01, Cultrex 83, Cultrex 87, or UltiMatrix 96 for 5 days as determined by levels of intracellular ATP (Cell Titer Glo - CTG). Data represent mean ± SD. Statistical significance was determined by 1-way ANOVA. \*\*\*\**P* ≤ 0.0001. (B) Growth of mouse organoids mT69A and mT69B cultured in Matrigel 04, Matrigel 01, Cultrex 83, Cultrex 87, or UltiMatrix 96 for 3 days as determined by levels of intracellular ATP (CTG). Data represent mean ± SD. Statistical significance was determined by 1-way ANOVA. \*\*\*\**P* ≤ 0.0001. (C) Dose- response curves for hF24 treated with gemcitabine, paclitaxel, SN-38, oxaliplatin, 5-FU, and trametinib during culture in Matrigel 04, Matrigel 01, Cultrex 83, Cultrex 87, or UltiMatrix 96. Data represent mean ± SD of triplicate values fitted with a four-parameter log-logistic function. (D) Dose-response curves for mT69B treated with gemcitabine, paclitaxel, SN-38, oxaliplatin, 5-FU, and trametinib during culture in Matrigel 04, Matrigel 01, Cultrex 83, Cultrex 87, or UltiMatrix 96. Data represent mean ± SD of triplicate values fitted with a four-parameter log-logistic function.

### BME has minimal impact on drug response

To investigate the influence of BME on drug response, we measured the dose-response of both mouse and human organoids to standard-of-care chemotherapies: gemcitabine, paclitaxel, SN-38 (active metabolite of irinotecan), oxaliplatin, and 5-fluorouracil (5-FU), as well as the targeted small molecule trametinib (third generation MEKi) (**Supplementary Figure 1**). Organoids were allowed to reform and then subjected to pharmacotyping. We found that the dose- response curves for gemcitabine, SN-38, and trametinib across all BMEs were largely overlapping in both hF24 and hM1F (n=3/3) (**Figure 1C-D**, **Supplementary Figure 3A-B**). The dose response curves in hT1 were also consistent across BMEs for SN-38 (n=2/2), but exhibited some variability for gemcitabine (n=1/2) and trametinib (n=1/2) between Matrigel, Cultrex, and Ultimatrix **(Supplementary Figure 3C**). Paclitaxel dose-response curves across all BMEs were also largely overlapping in hM1F (n=3/3) (**Figure 1D**), hT1 (n=2/2) **(Supplementary Figure 3C**), and hF24 (n=2/3) (**Figure 1C, Supplementary Figure 3A).** Oxaliplatin also displayed consistent dose- response curves regardless of BME hF24 (n=3/3), hM1F (n=2/3), and hT1 (n=2/2) (**Figure 1C-D**, **Supplementary Figure 3A-C**). Additionally, 5-FU exhibited overlapping dose-response curves in hF24 (n=3/3) (**Figure 1C**, **Supplementary Figure 3A**) and hM1F (n=2/3) (**Figure 1D**, **Supplementary Figure 3B**). However, we observed more variability in dose-response curves for 5-FU in hT1 as a function of BME in both experimental repeats conducted (**Supplementary Figure 3C**).

Mouse organoids were less sensitive to all agents tested relative to hPDOs, with oxaliplatin and 5-FU dose-response curves failing to reach 50% cytotoxicity (**Supplementary Figure 3D-E**). Gemcitabine and SN-38 mouse organoid dose response curves exhibited no consistent or substantial shifts when cultured in different BMEs (**Supplementary Figure 3D-E**). Dose-response curves for trametinib were also largely overlapping regardless of BME for both mT69A (n=3/4) and mT69B (n=3/4), although both organoid lines were more resistant in 1 out of 4 experimental replicates when cultured in UltiMatrix 96 (**Supplementary Figure 3D-E**). We observed a similar dose response shift for paclitaxel when mouse organoids were cultured in Ultimatrix (n=1/4) (**Supplementary Figure 3D-E**). Overall, these data demonstrate that choice of BME does not induce consistent shifts in dose-response curves for PDOs treated with standard-of-care chemotherapies or trametinib, suggesting that either Matrigel, Cultrex, or UltiMatrix would be appropriate for PDO pharmacotyping analyses.

### BME does not induce significant alterations in dose-response parameters

We next compared dose-response parameters to gain further insight into subtle changes in dose response curves both within individual experiments and across all experimental replicates. Within each individual experimental replicate, we performed statistical analyses to identify differences in the half maximum inhibitory concentration (IC50s), Hill slope, area under the curve (AUC), and standard deviation of the residuals (Sy.x) between BME lots and sources (*22*).

We observed a modest number of instances in which IC50s were statistically significant between BMEs within experimental replicates (**Figure 2A-B, Supplementary Figure 4**). Significant pairwise comparisons in IC50 were predominantly observed between Matrigel and Cultrex or between Matrigel and UltiMatrix. Across all organoids, drug treatments, and BME comparisons, statistical differences between lots of Matrigel (04 vs 01) were observed only once in a single experimental replicate (Paclitaxel, hF24) (**Supplementary Figure 4A**). Similarly, statistical differences between lots of Cultrex (83 vs 87) were observed in only two experimental replicate (SN-38, hM1F; 5-FU, mT69A) (**Supplementary Figure 4B**). These data indicate that lot within the same BME source has little impact on drug IC50. Differences in IC50s for oxaliplatin and trametinib were rarely significant, occurring in 1/16 experiments (hT1, Matrigel 04 vs. Cultrex 87) (**Supplementary Figure 4C**) and 2/16 experiments (hM1F, Matrigel 01 vs. UltiMatrix; mT69A, Matrigel 01 vs. UltiMatrix, Cultrex 87, Cultrex 83 vs. UltiMatrix) (**Supplementary Figure 4B, D**), respectively. Conversely, we consistently observed statistical differences in gemcitabine IC50 between Matrigel and Cultrex/UltiMatrix in each experimental replicate of hF24 and hM1F (**Figure 2**, **Supplementary Figure 4A-B**), as well as in hT1 (n=1/2), mT69A (n=1/4), and mT69B (n=2/4) (**Supplementary Figure 4C-E**). Statistical differences in paclitaxel IC50 were observed in hF24 (n=1/3), hM1F (n=1/3), hT1 (n=2/2), and mT69B (n=1/4) (**Supplementary Figure 4A-C, E**). Lastly, differences in 5-FU IC50 were observed in only one experimental replicate for hF24, mT69A, and mT69B (**Figure 2A**, **Suppelementary Figure 4D-E**). To further compare differences in dose-response across experimental repeats, we calculated average IC50 concentrations (**Figure 2C-D**, **Supplementary Figure 6**). For all hPDOs, we observed no statistically significant alterations in mean IC50s over experimental repeats calculated for any drug as a function of BME (**Figure 2C-D**, **Supplementary Figure 6A**). Similarly, mouse organoids exhibited no statistically significant shifts in drug IC50, regardless of which BME organoids were plated in (**Supplementary Figure 6B-C**).

**Figure 2.**
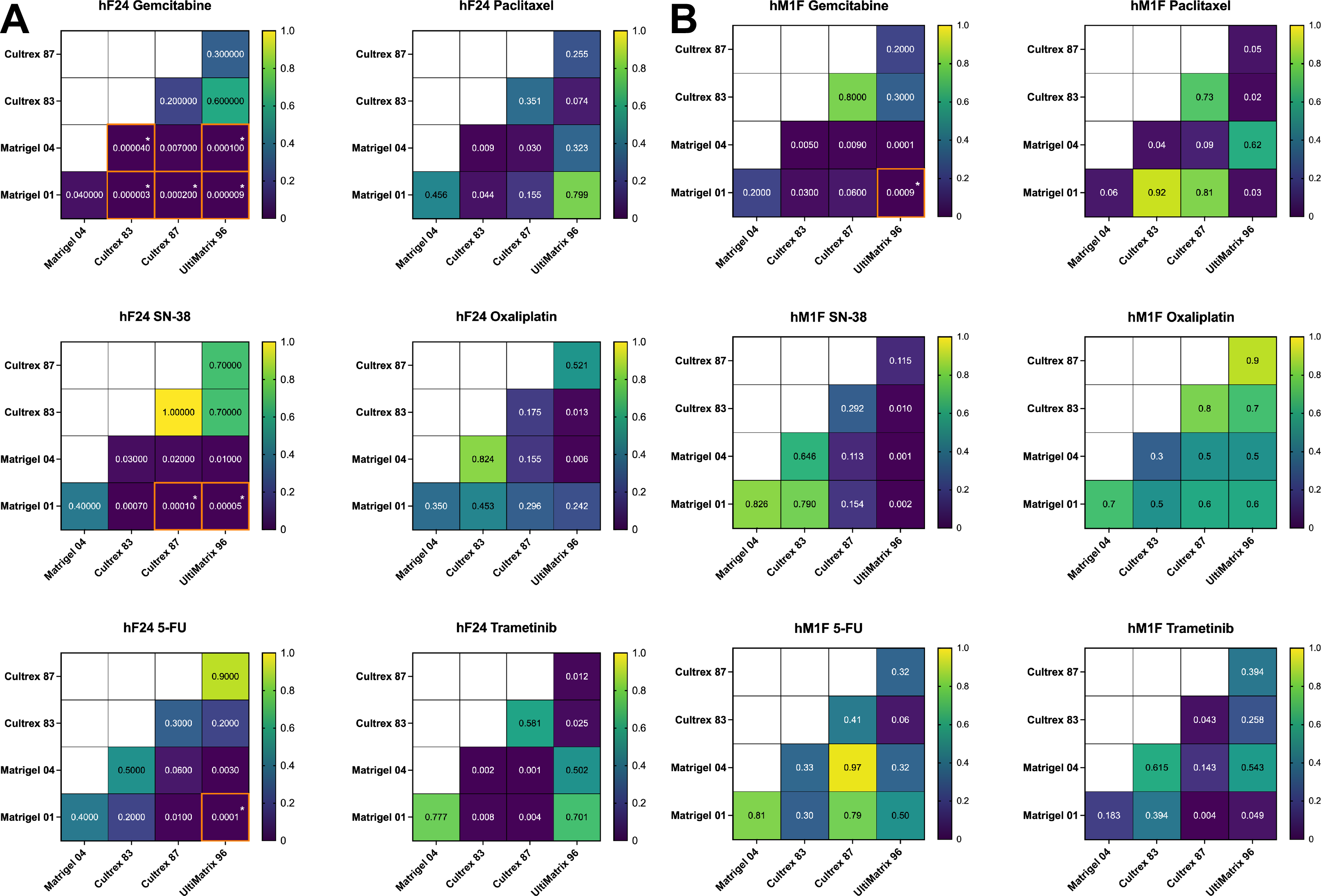
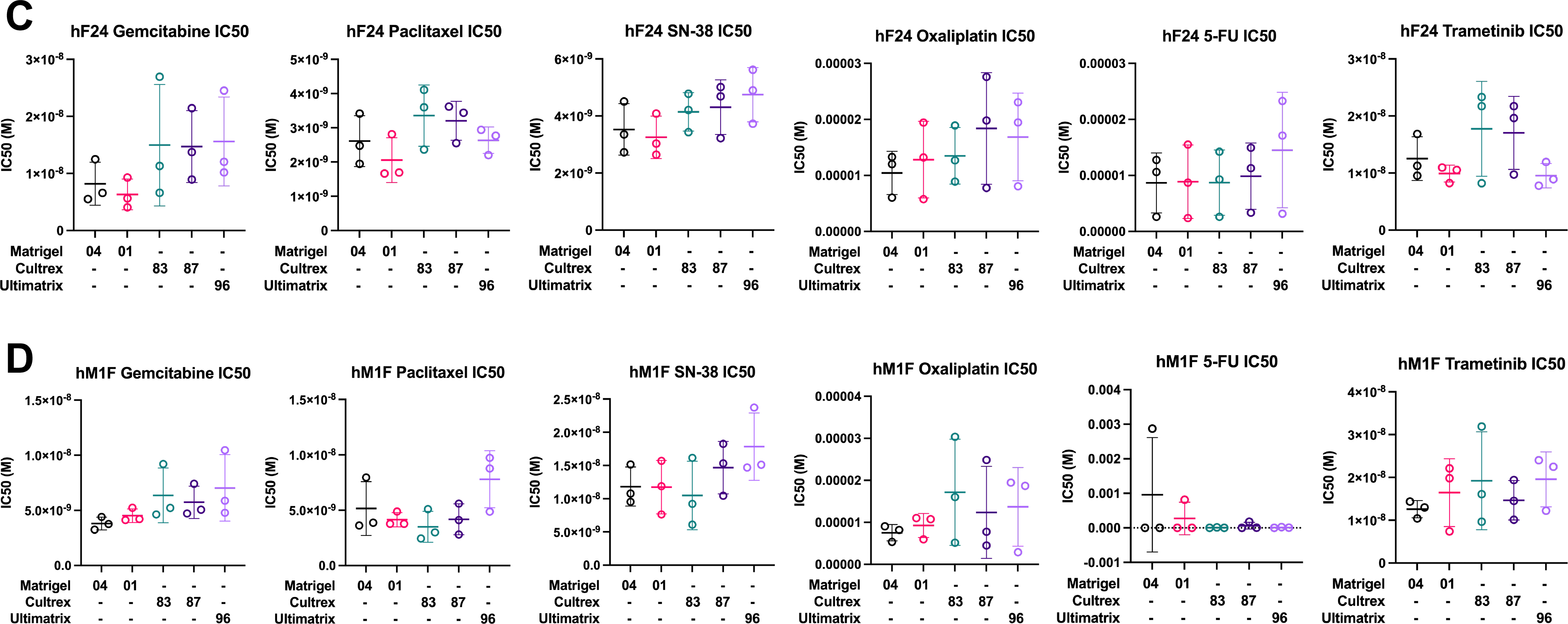
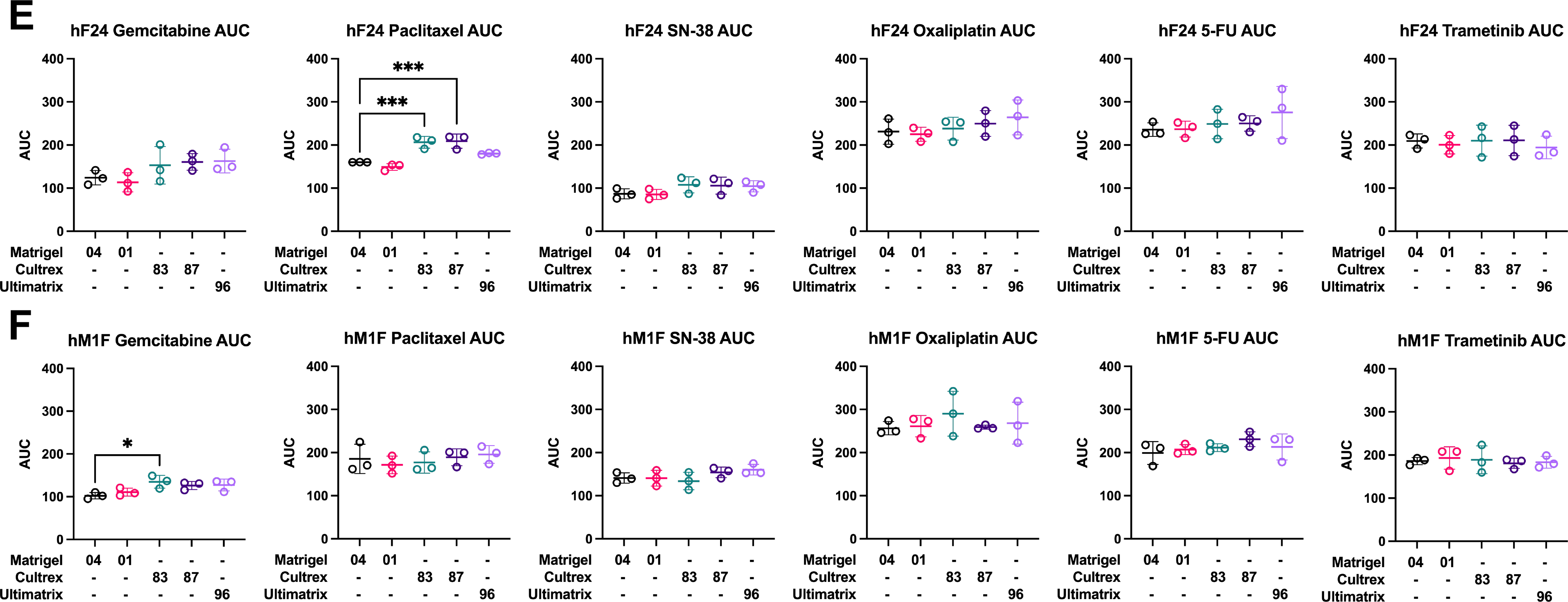
Basement membrane extracts exert minimal effects on drug IC50 and dose- response AUC. (**A-B**) *P* value heatmaps from pairwise comparisons of IC50 between BMEs for hF24 and hM1F. Each square represents pairwise t-tests. After Bonferroni adjustment, t-tests were considered significant if *P* < 0.00027778. Statistically significant comparisons are denoted by an asterisk and orange border. (**C-D**) Gemcitabine, paclitaxel, SN-38, oxaliplatin, 5-FU, and trametinib IC50 values (M) for hF24 and hM1F across different BMEs. Data represent mean ± SD of 3 experimental replicates. (**E-F**) Gemcitabine, paclitaxel, SN-38, oxaliplatin, 5-FU, and trametinib AUC values for hF24 and hM1F across the different BMEs. Data represent mean ± SD of 3 experimental replicates. Statistical significance was determined by 1-way ANOVA. \**P* ≤ 0.05, \*\*\**P* ≤ 0.001 vs. Matrigel 04.

Hill slope is another metric of drug potency that has been suggested to offer higher predictive power than IC50 (*23*). Pairwise comparisons of Hill slope revealed little to no significant differences among BMEs on a per-drug or per-organoid basis (**Supplementary Figure 5**). The only instances in which Hill slopes were statistically distinct between BMEs were in mouse organoids treated with 5-FU, likely due to the resistant nature of these organoids to 5-FU yielding disparate dose-response curves (**Supplementary Figure 5D-E**).

We also rarely observed changes in average dose-response curve AUCs across BMEs. Between BME lot, there were not significant AUC differences. AUCs for hF24 treated with paclitaxel were slightly (1.28- to 1.3-fold) but significantly higher in Cultrex 83 or Cultrex 87 compared to Matrigel 04 (**Figure 2E**). Similarly, calculated AUCs for hM1F treated with gemcitabine were 1.3-fold higher on average in Cultrex 83 compared to Matrigel 04 (**Figure 2F**). Lastly, AUCs for mT69A treated with paclitaxel were increased in Ultimatrix 96 compared to Matrigel 04 (1.28-fold) (**Supplementary Figure 7B**). In all other mouse and human drug-organoid combinations tested, there were not significant differences in mean AUC across multiple experiments. These data suggest that while minimal differences in drug response exist, the most rigorous approach is to complete pharmacotyping analyses using the same commercial BME source.

We next sought to analyze dose-response curve goodness-of-fit as a parameter of plating consistency in each BME within and across experimental replicates. We measured Sy.x as an estimate of goodness-of-fit. In general, higher Sy.x values indicate larger deviations of data points from the fit regression curve. Sy.x values for Cultrex 83 were relatively consistent across experimental replicates, with low absolute values (<10%) and low inter-experimental variability (**Figure 3A-B**). The largest variations in Sy.x across experimental replicates in Cultrex were hF24 treated with SN-38 and 5-FU, and hM1F treated with gemcitabine and oxaliplatin (**Figure 3A-B**). Sy.x values were larger on average for Cultrex 83 in hM1F treated with oxaliplatin (mean = 20.37%). Cultrex 87 performed consistently as well (ranging from 1.6 – 11.5%), with the most statistical variance across experiments observed in hF24 treated with gemcitabine (**Figure 3A**). UltiMatrix 96, relative to the other BMEs, exhibited higher experimental variability in Sy.x values, with greater variance observed in hF24 treated with gemcitabine (4.1 – 11%) and oxaliplatin (7.6 – 22.3%); and hM1F treated with gemcitabine (4.0 – 15.8%), paclitaxel (4.6 – 14.1%), oxaliplatin (7.5 – 26%), and trametinib (4.8 – 12.6%) (**Figure 3A-B**). Matrigel 04 yielded consistent and relatively low Sy.x values (<10%) across all experimental replicates, with the exception of hF24 treated with 5-FU (6.3 – 13.3%) (**Figure 3A**). Similarly, Matrigel 01 demonstrated low and consistent Sy.x values across treatments, experimental replicates, and hPDOs tested, as evidenced by low standard deviations and low Sy.x absolute values **(Figure 3A, Supplementary Figure 8**). Overall, these results indicate better dose-response curve goodness-of-fit from experiment to experiment when human organoids are cultured in Matrigel.

**Figure 3.**
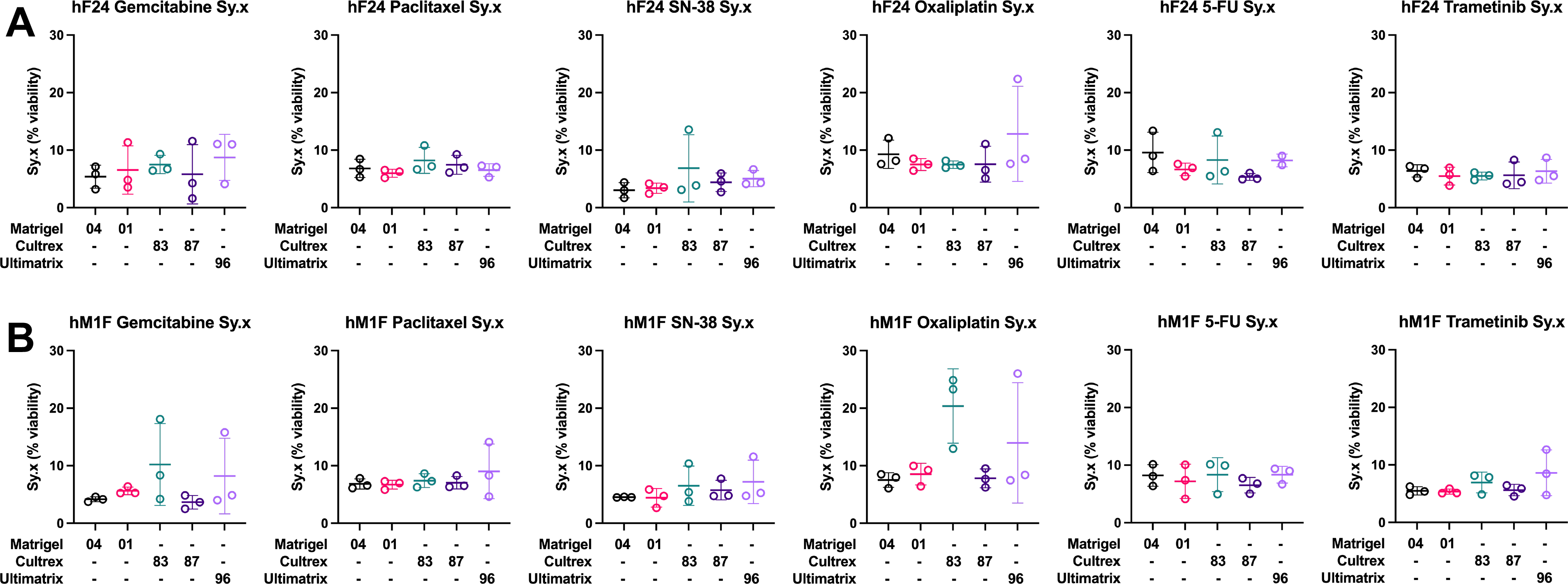
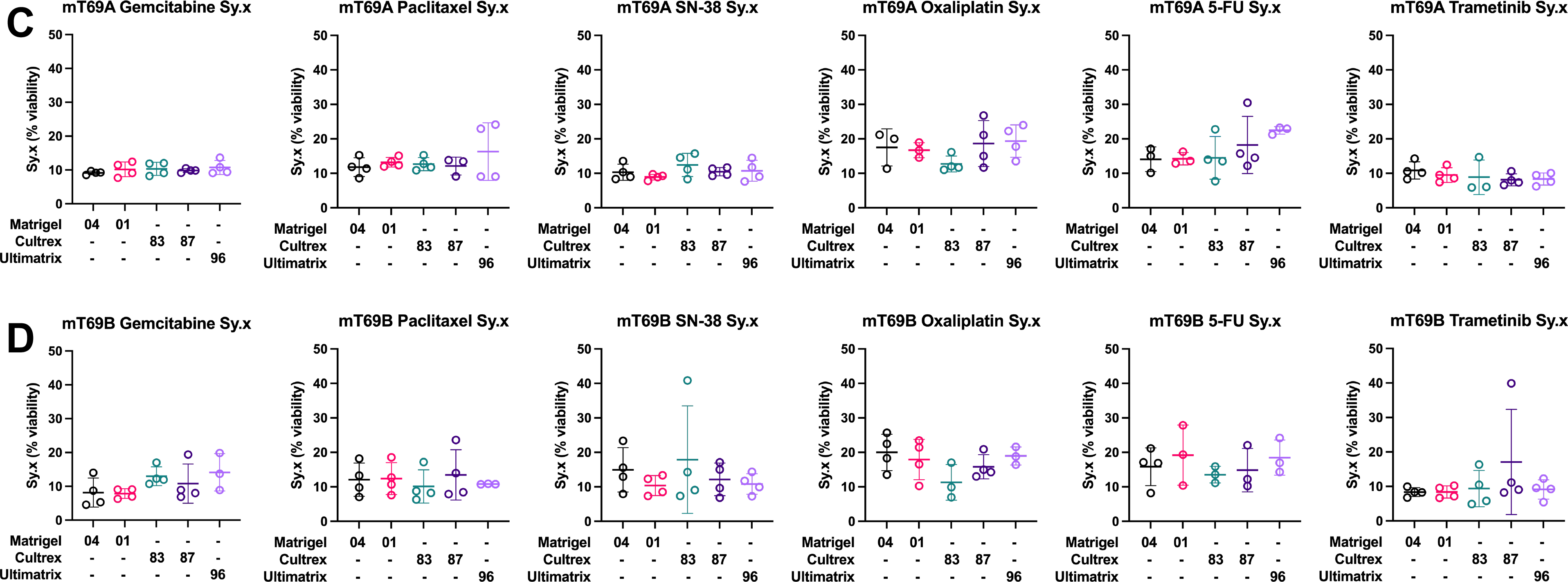
Dose-response curve goodness of fit is not significantly altered by basement membrane extract. (**A-B**) Standard deviation of the residuals (Sy.x) for hF24 and hM1F treated with gemcitabine, paclitaxel, SN-38, oxaliplatin, 5-FU, and trametinib across the different BMEs. Data represent mean ± SD of 3 experimental replicates. (**C-D**) Sy.x for mT69A and mT69B treated with gemcitabine, paclitaxel, SN-38, oxaliplatin, 5-fluorourical (5-FU), and trametinib across the different BMEs. Data represent mean ± SD of 4 experimental replicates. Outliers were determined using Grubbs’ test α = 0.05.

For mouse organoids, similar trends were observed, albeit with higher absolute Sy.x values across almost all conditions (range: 4.5 – 40.8%) (**Figure 3C-D**). In general, mouse organoids are more sensitive to single cell dissociation protocols and have higher growth rates, both of which contribute to increased variance and necessitate shorter pharmacotyping experimental time courses (3 days for mouse organoids vs 5 days for hPDOs). Over the course of 4 experimental replicates, UltiMatrix 96 demonstrated a high degree of statistical variability in Sy.x values relative to the other BMEs tested, notably for mT69A treated with paclitaxel (9.0 - 24.1%) and oxaliplatin (13.5 - 23.9%), and mT69B treated with gemcitabine (8.81 - 19.8%) (**Figure 3C-D**). Relative to the other BMEs tested, Cultrex lots also exhibited high levels of variability in Sy.x. Cultrex 83 exhibited a high degree of variability for mT69A treated with 5-FU (7.7 - 22.7%) and trametinib (5.9 - 14.7%), and mT69B treated with paclitaxel (6.3 - 17.1%) and SN-38 (7.3 - 40.8%) (**Figure 3C-D**). Similar variability in Sy.x across experimental replicates was observed for Cultrex 87 in mT69A treated with oxaliplatin (11.7 - 26.8%) and 5-FU (12.2 - 30.5%), and mT69B treated with gemcitabine (6.9 - 19.4%), paclitaxel (7.9 - 23.6%), and trametinib (8.2 - 39.9%) (**Figure 3C-D**). Sy.x values were consistent for mT69B treated with paclitaxel when cultured in Ultimatrix 96 (10.7 - 10.9%) (**Figure 3D**). Both Matrigel 04 and Matrigel 01 showed relatively consistent Sy.x values in the mouse organoid context but exhibited some biological and experimental variance. Most notably, mT69A and mT69B are relatively resistant to oxaliplatin and 5-FU and exhibited relatively high Sy.x absolute values (20-30%) in these conditions compared other drug treatments (**Figure 3C-D**).

Overall, we observed no statistically significant differences in average Sy.x values for both human and mouse organoids across BMEs and all drugs (**Figure 3, Supplementary Figure 8**). However, for both human and mouse organoids, both lots of Matrigel exhibited more consistent Sy.x values across experimental replicates compared to Cultrex or UltiMatrix, as evidenced by lower variance in Sy.x values across experiments (**Figure 3**). Lot-to-lot variability was more pronounced for Cultrex compared to Matrigel for both human and mouse organoids. Collectively, these data indicate that drug dose-response curve fitting is stable across the BMEs tested.

### Gene expression and organoid subtype classification are not influenced by BME

Although we did not observe substantial shifts in drug response as a function of BME lot or source, alterations in intracellular signaling and cell state represented another potential variable influenced by BME composition. Therefore, we also investigated BME induced changes in gene expression or PDA subtype classification (*24*). To this end, mouse organoids (mT69A, mT69B, mT9) and human PDOs (hF24, hM1F, hT1) were each cultured in each BME (Matrigel 04, Matrigel 01, Cultrex 83, Cultrex 87, and Ultimatrix 96) for 3 days and subjected to RNA-seq analyses (**Supplementary Figure 1**). Multivariate analysis of resultant Z-scores of organoid normalized counts indicated that both mouse and human samples clustered well based on organoid line (**Figure 4A**, **Supplementary Figure 9C-D**). However, samples did not cluster significantly based on BME (**Figure 4A, Supplementary Figure 9C-D, F-G**). Gene expression was largely unchanged between Matrigel lots in both human and mouse organoids (0-2 differentially expressed genes, 20 in hM1F) (**Supplementary Figure 9A-B**). Similarly, minimal gene expression changes were observed in human and mouse organoids between Cultrex lots (0-4 differentially expressed genes) (**Supplementary Figure 9A-B**). Between BME sources, we observed more changes in gene expression, but these were not consistent between organoid lines. For example, hT1 (Stage 2B) displayed 522 differentially expressed genes between Matrigel 04 and UltiMatrix, whereas hF24 (Stage 4) and hM1F (Stage 4, lung metastasis) presented 0 (**Supplementary Figure 9A**). Similarly, mT69B (KPC PDA primary tumor) displayed 90 differentially expressed genes between Matrigel 04 and Cultrex 87, whereas mT69A (KPC PDA metastasis) presented with 13 (**Supplementary Figure 9B**). These results indicate that BME does not substantially alter PDO gene expression profiles.

**FIgure 4.**
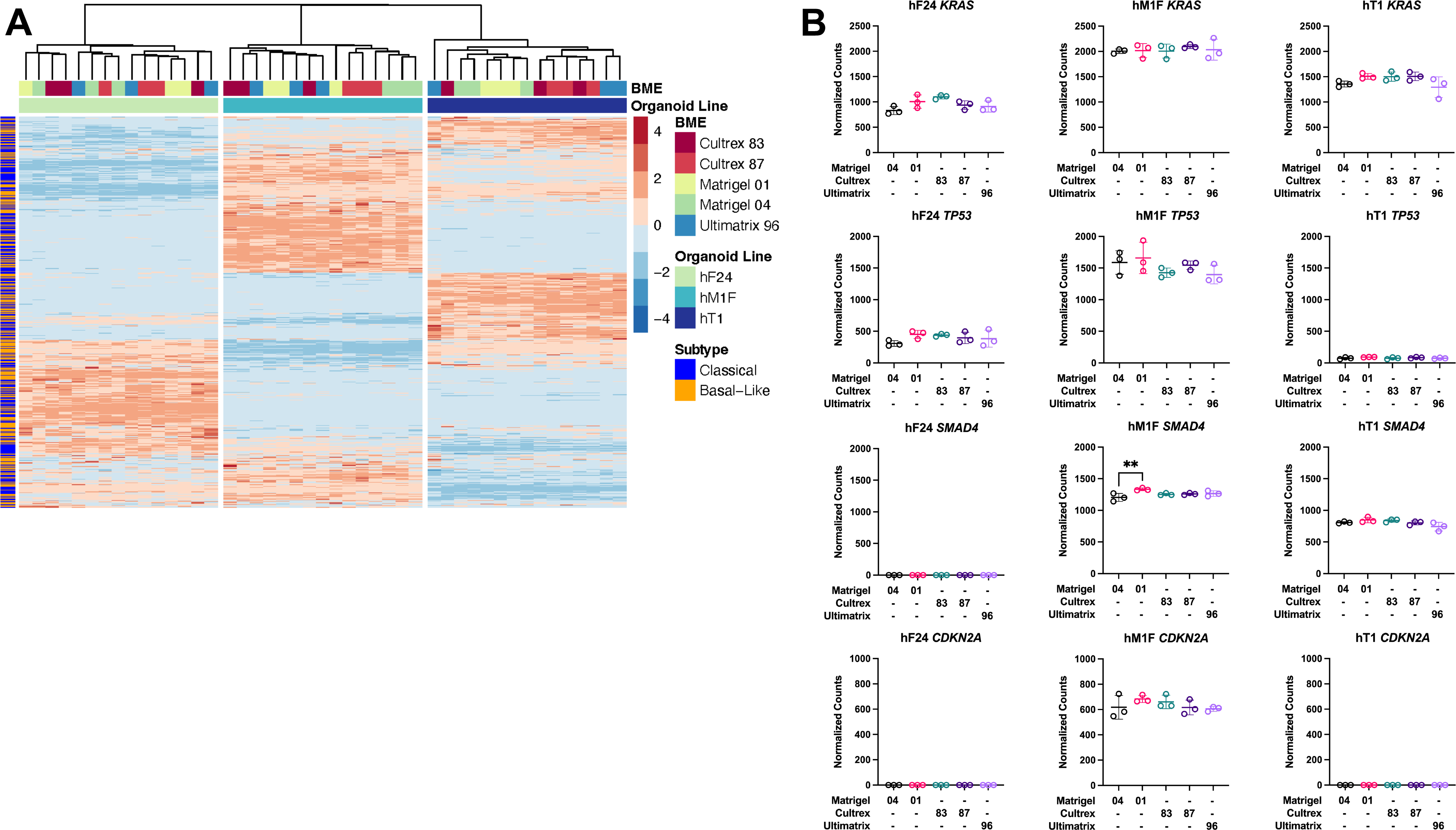
Human patient-derived organoid gene expression profiles are consistent across basement membrane extract types. (**A**) Expression heatmap of Z-scores of normalized counts of classical or basal-like hallmark genes in hF24, hM1F, and hT1 when cultured in different BMEs. (B) Normalized counts for hallmark PDA genes *KRAS, TP53, SMAD4,* and *CDKN2A*. Data represent mean ± SD of 3 technical replicates. Statistical significance was determined by 1-way ANOVA. \*\**P* ≤ 0.01 vs. Matrigel 04.

Previous work defined molecular subtypes of PDA. Patients with basal-like PDA exhibit poor prognoses and increased therapeutic resistance compared to the classical subtype (*24*). Changes to molecular subtype would have major implications on the predictive power of PDOs in precision medicine approaches. To determine whether BME influenced hPDO subtype classification, we delineated significant changes to enrichment of basal-like and classical differentially expressed genes (see **Supplementary File** for gene list). Pairwise comparisons of subtype gene enrichment indicated no significant difference in basal-like or classical gene enrichment between Matrigel 04 and Matrigel 01, or between Cultrex 83 and Cultrex 87 (**Supplementary Figure 9E**). Notably, a number of comparisons that were enriched for one hallmark type were also enriched for the other (**Supplementary Figure 9E**). For instance, hM1F displayed upregulation of both basal-like as well as classical hallmark genes when cultured in Cultrex 83, Cultrex 87, or UltiMatrix **(Supplementary Figure 9E**). Indeed, overall, there was a greater enrichment for the classical subtype signature, in concordance with previous work demonstrating that organoid models primarily exhibit RNA states reminiscent of the classical subtype (*16*). Analysis of individual genes implicated in the etiology of PDA revealed that BME did not significantly alter hPDO expression levels of *KRAS*, *TP53*, or *CDKN2A.* A slight but statistically significant increase in *SMAD4* expression was detected in hM1F cultured in Matrigel 01 compared to Matrigel 04 (**Figure 4B**). Similarly, we observed slight yet statistically significant increases in *ERBB3* in all three hPDOs cultured in Cultrex or UltiMatrix (**Supplementary Figure 9H**). Expression of *EGFR* was increased in hT1 cultured in Cultrex 87 relative to Matrigel 04 (**Supplementary Figure 9H**). However, we did not observe any significant changes in expression levels of *MYC*, *GATA6*, *MAP2K1*, *MAP2K2*, *SOX9*, *RB1*, *BRCA2*, and *ERBB2*, as well as genes related to drug resistance such as *HNF1A* and *ABCB1* (**Supplementary Figure 9H**). Overall, these data indicate that the potential influence of BME on organoid gene expression, cell state, and subsequent response to targeted therapies, must be carefully considered when developing protocols for organoid-guided therapy studies.

## Discussion

The ability of PDOs to mirror patient genetics, transcriptomics, and drug response in retrospective studies prompted the onset of several clinical trials to test the benefit of using PDOs to inform treatment decisions. However, previous studies demonstrated the plasticity of PDOs in response to varying culture conditions and potential impact on drug response (*16, 17*). With multi-institutional clinical trials underway, the need to investigate factors that impact the robustness of PDO genetics, transcriptomics, and drug testing is imperative. Basement membrane extracts are central to most PDO efforts. Previous studies demonstrated variability in protein composition, presence of xenogenic bioactive compounds, and scaffold stiffness between batches. Despite this variability, commercially available BMEs remain critical and prevalent in organoid efforts. Given the multi-year nature of PDO-related clinical trials, more than one lot of BME will be required for the duration of these studies. In addition, limitations in availability due to supply chain issues has necessitated that some efforts switch between alternate commercial sources. However, the influence of BME source and lot on organoid response to drug treatment was unknown.

To investigate the extent to which BME impacts growth, drug response, and transcriptomics, we compared various commercially available BMEs on previously established mouse and human PDA organoids. We consistently observed a decrease in mouse and human organoid growth when cultured in Cultrex or UltiMatrix compared to Matrigel. While this could potentially arise from a lack of an adaptation period, multiple passages of mouse organoids in Cultrex does not rescue their growth rate. Despite this growth deficit, we observed no substantial shifts in drug response or gene expression as a function of BME source or lot. The generation efficiency of PDA organoids in Matrigel is 75% (*8*). This study made use of previously established organoids. Given the impact of BME source on PDO growth, it will be important to delineate the effect of BME on isolation efficiency in future work. For established organoids, Matrigel, Cultrex, or UltiMatrix BMEs are all viable options for pharmacotyping experiments. Altogether, this work demonstrates that precision medicine approaches using organoids are robust, and that ongoing clinical trials are unlikely to be affected by different lots or commercial sources of BME.

## Methods

### Organoids and cell culture conditions

Organoids used in this study were generated at Cold Spring Harbor Laboratory (*7, 8, 25*). Human patient-derived and mouse organoids were cultured as described. In short, mouse organoid media contained Advanced DMEM/F12, HEPES, Glutamax, Penicillin/Streptomycin, EGF, FGF-10, Gastrin, A83-01, B27 supplement, *N*-acetylcysteine, nicotinamide, Noggin, and R- spondin1. For human organoid culture, Wnt3a was also included in the media. Organoid nomenclature is defined as: hT – human tumor obtained from resection; hF – human fine-needle biopsy obtained by fine-needle aspiration or core biopsy; hM – human metastasis obtained from direct resection of metastases following rapid autopsy or VATS resection; mT – murine tumor. All organoid models were routinely tested for *Mycoplasma* at Salk Institute. Additional hPDO and mouse organoid characteristics are available in **Table S1** and **Table S2**.

### Basement membrane extracts

Both human and mouse organoids were maintained in Corning Matrigel Lot #1062004 during the growth and expansion phase. For pharmacotyping comparisons, organoids were plated in either Corning Matrigel (Growth Factor Reduced, Phenol Red Free) Lot #1062004 (Matrigel 04), Corning Matrigel (Growth Factor Reduced, Phenol Red Free) Lot #0287001 (Matrigel 01), RnD Cultrex Reduced Growth Factor BME Lot #1564183 (Cultrex 83), RnD Cultrex Reduced Growth Factor BME Lot #1586187 (Cultrex 87), or Cultrex UltiMatrix Reduced Growth Factor BME Lot #1637796 (UltiMatrix 96).

### Pharmacotyping of organoids

Organoids were dissociated into single cells using a diluted solution of TrypLE Express (Thermo Fisher). Equal numbers of single cells (1,500 viable cells per well) were plated in a 20 uL suspension of 10% BME in ultra-low attachment 384 well plates (Corning). Twenty-four hours after plating, reformation of organoids was visually verified and therapeutic compounds were applied using a digital dispenser (Tecan). Chemotherapies were tested in triplicate 10-point dose-response curves. Gemcitabine, paclitaxel, SN-38 (active metabolite of irinotecan), and trametinib ranged from 0.5nM – 5μM; oxaliplatin and 5-fluorouracil ranged from 50nM – 100μM. Compounds were dissolved in DMSO, and all treatments were normalized to 0.5% DMSO content. Mouse organoids underwent treatment for 3 days, while human organoids underwent treatment for 5 days. After treatment, cell viability was assessed using Cell Titer Glo (Promega) per manufacturer’s instructions on a Tecan Spark Cyto plate reader. A four-parameter log-logistic function with upper limit equal to the mean of the lowest dose values was fit to the data (viability vs. dose) with Graphpad Prism 9. The half maximal inhibitory concentration (IC50) values, area under the curve (AUC), and standard deviation of the residuals (Sy.x) for each dose-response curve were calculated using Graphpad Prism 9. Each pharmacotyping experiment was carried out at least two times.

### RNA isolation

For RNA-seq comparisons, organoids were maintained in Corning Matrigel Lot #1062004 (Matrigel 04) and then plated in either Corning Matrigel (Growth Factor Reduced, Phenol Red Free) Lot #1062004 (Matrigel 04), Corning Matrigel (Growth Factor Reduced, Phenol Red Free) Lot #0287001 (Matrigel 01), RnD Cultrex Reduced Growth Factor BME Lot #1564183 (Cultrex 83), RnD Cultrex Reduced Growth Factor BME Lot #1586187 (Cultrex 87), or Cultrex UltiMatrix Reduced Growth Factor BME Lot #1637796 (Ultimatrix 96). Each BME condition was plated in triplicate, and organoids were grown for 3 days before harvesting in TRIzol (Life Technologies) and snap frozen. RNA was isolated using the RNeasy Mini Kit (Qiagen). RNA quality control was performed for all samples using a Qubit Fluorometer (Life Technologies) and TapeStation (RIN greater than 8.5) (Agilent) before RNA-seq analyses.

### RNA-seq analyses

Library construction was conducted using the Truseq RNA library prep kit (Illumina) with 500ng RNA input. Raw reads were trimmed with Trim Galore v0.4.4_dev (https://www.bioinformatics.babraham.ac.uk/projects/trim_galore/) and quality-checked with FastQC v0.11.8 (http://www.bioinformatics.babraham.ac.uk/projects/fastqc). Trimmed reads were then aligned to the hg38 human reference genome with STAR aligner v2.5.3a (*26*), and converted to gene counts with HOMER’s analyzeRepeats.pl script (*27*).

### Differential gene expression analysis

Gene counts where normalized and queried for differential expression using DESeq2 v1.30.0 (*28*). For each pairwise comparison, genes with fewer than 10 total raw counts across all samples were discarded prior to normalization, and genes with an absolute log2foldchange > 1 and an FDR-corrected p-value ≤ 0.05 were pulled as significant.

### Statistics

A four-parameter log-logistic function with upper limit equal to the mean of the lowest dose values was fit to the dose-response data (viability vs. dose) with Graphpad Prism 9. The half maximal inhibitory concentration (IC50) values, area under the curve (AUC), and standard deviation of the residuals (Sy.x) for each dose-response curve were calculated using Graphpad Prism 9. Each pharmacotyping experiment was carried out for n = 2-4 experimental replicates. Differences among means were analyzed using 1-way ANOVA with Dunnett’s multiple comparisons test. Outliers were determined using Grubbs’ test α = 0.05.

To compare estimated dose-response curve parameters across different BMEs within an experimental replicate, t-tests were conducted on the IC50 and Hill slope to test the null hypothesis of zero difference in the parameters among BMEs using R 4.0.3 and package drc (*22, 29*). The Bonferroni method was used to adjust for multiple comparisons, and a comparison between a pair of BMEs was considered significant if the *P* value was below 0.05 divided by the total number of pairwise comparisons. Consequently, pairwise t-tests were considered significant if *P* < 0.00027778 (hPDO) or *P* < 0.00041667 (mouse organoids).

To assess the model fitting and compare the log-logistic models with other alternatives, including Weibull models, a sensitivity analysis was conducted. The Akaike Information Criterion (AIC) was used for the comparison. The four-parameter log-logistic (LL4) model consistently yielded smaller AIC values, suggesting this model as the most appropriate model-fitting strategy. Analyses presented from the aforementioned t-tests on IC50 and Hill slope were conducted first on data fitted with the LL4 model, then on data fitted with Weibull models when appropriate.

## Author contributions

JL, SO, and KP maintained organoid cultures, acquired materials, and designed and conducted the experiments. JL and XL acquired and analyzed the data. KL performed statistical analysis on RNA-seq data. JZ directed and provided statistical analysis for dose-response curve comparisons. JL, XL, JZ, and DE wrote the manuscript. DE conceived the project and directed the experiments.

## Acknowledgements

The authors are supported by National Institutes of Health, National Cancer Institute R00CA024725 (Engle), P30DK120515 (Engle), T32CA009370 (Lumibao); the Pancreatic Cancer Action Network 19-20-ENGL (Engle), American Association for Cancer Research and the Lustgarten Foundation 21-20-67-ENGL (Engle), the University of California Tobacco-Related Disease Research Program T31KT1898 (Engle), and Padres Pedal the Cause/C3 #PTC2020 (Engle). We are grateful to the Rose Hills Foundation, Mission Cure Capital, LLC, the Mark Foundation for Cancer Research, and the Emerald Foundation, Inc. for supporting Dr. Engle. Additionally, this work was supported by the Razavi Newman Integrative Genomics and Bioinformatics Core Facility of the Salk Institute with funding from NIH-NCI CCSG: P30 014195, and the Helmsley Trust; the NGS Core Facility of the Salk Institute with funding from NIH-NCI CCSG: P30 014195, the Chapman Foundation and the Helmsley Charitable Trust; and the Stem Cell Core Facility of the Salk Institute with funding from the Helmsley Charitable Trust.

---

**Supplementary Figure 1.**
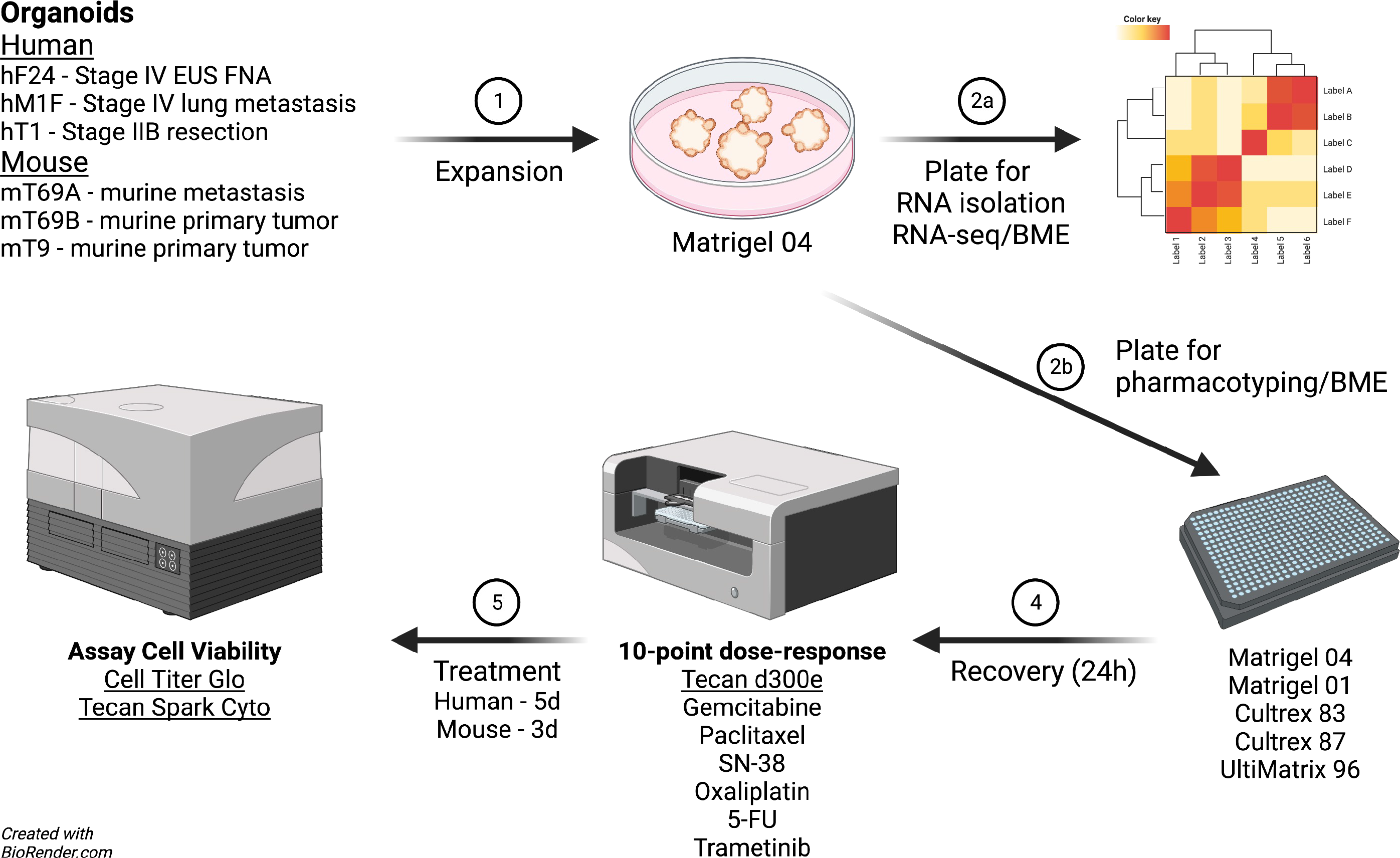
Experimental outline. Three biological replicates of human patient-derived organoids and mouse organoids were selected based on primary sample type and disease stage. Each organoid line was expanded to sufficient quantities for experimentation in Matrigel 04. Organoids were then single-cell dissociated and either plated in triplicate in 6-well plates for RNA isolation and subsequent RNA-seq, or plated for pharmacotyping in five 384-well plates in Matrigel 04, Matrigel 01, Cultrex 83, Cultrex 87, or UltiMatrix 96. Organoids were allowed to recover overnight. Next, organoids were treated with drugs and incubated for 5 days (human organoids) or 3 days (mouse organoids), followed by Cell Titer Glo to assess organoid viability.

**Supplementary Figure 2.**
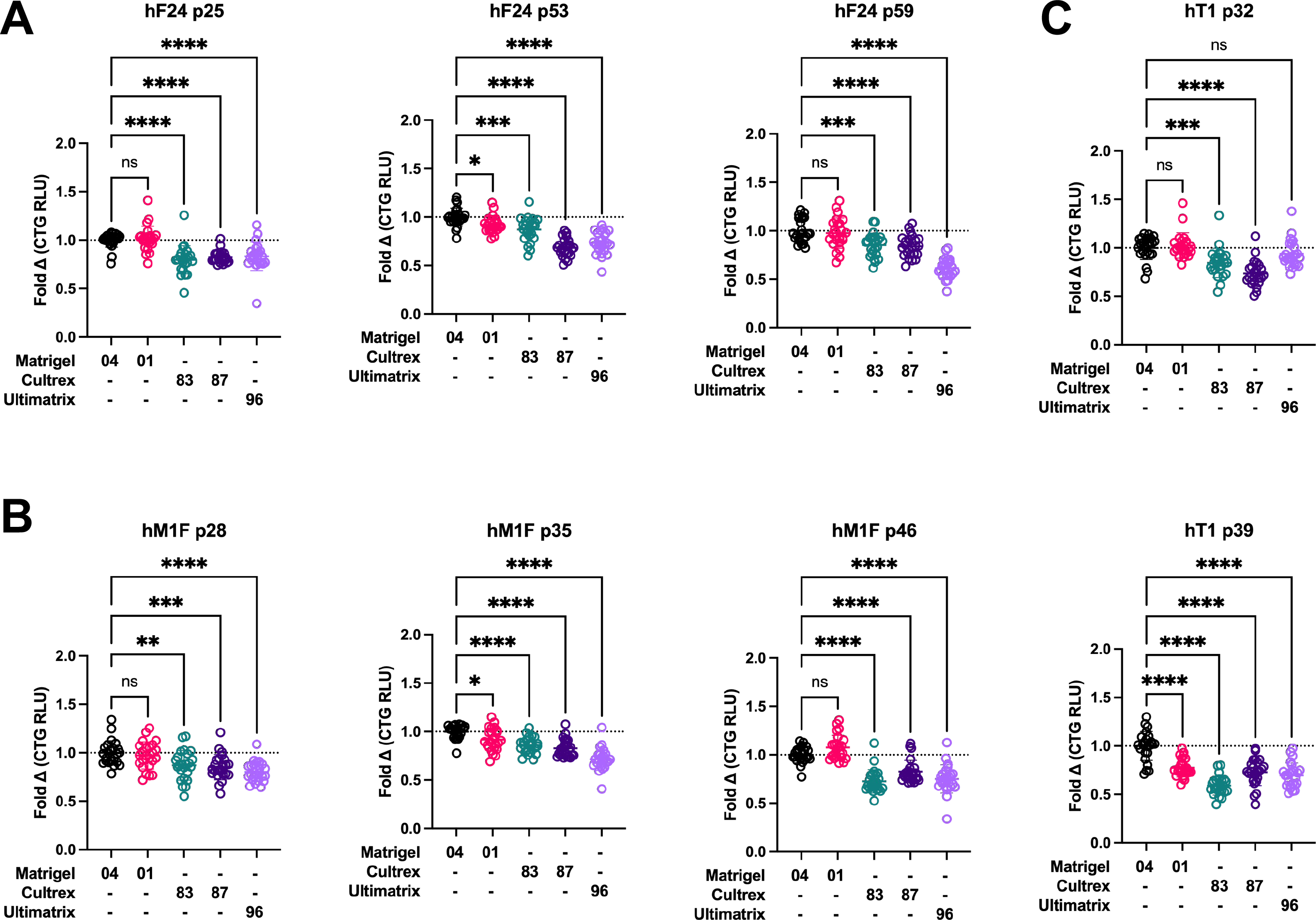
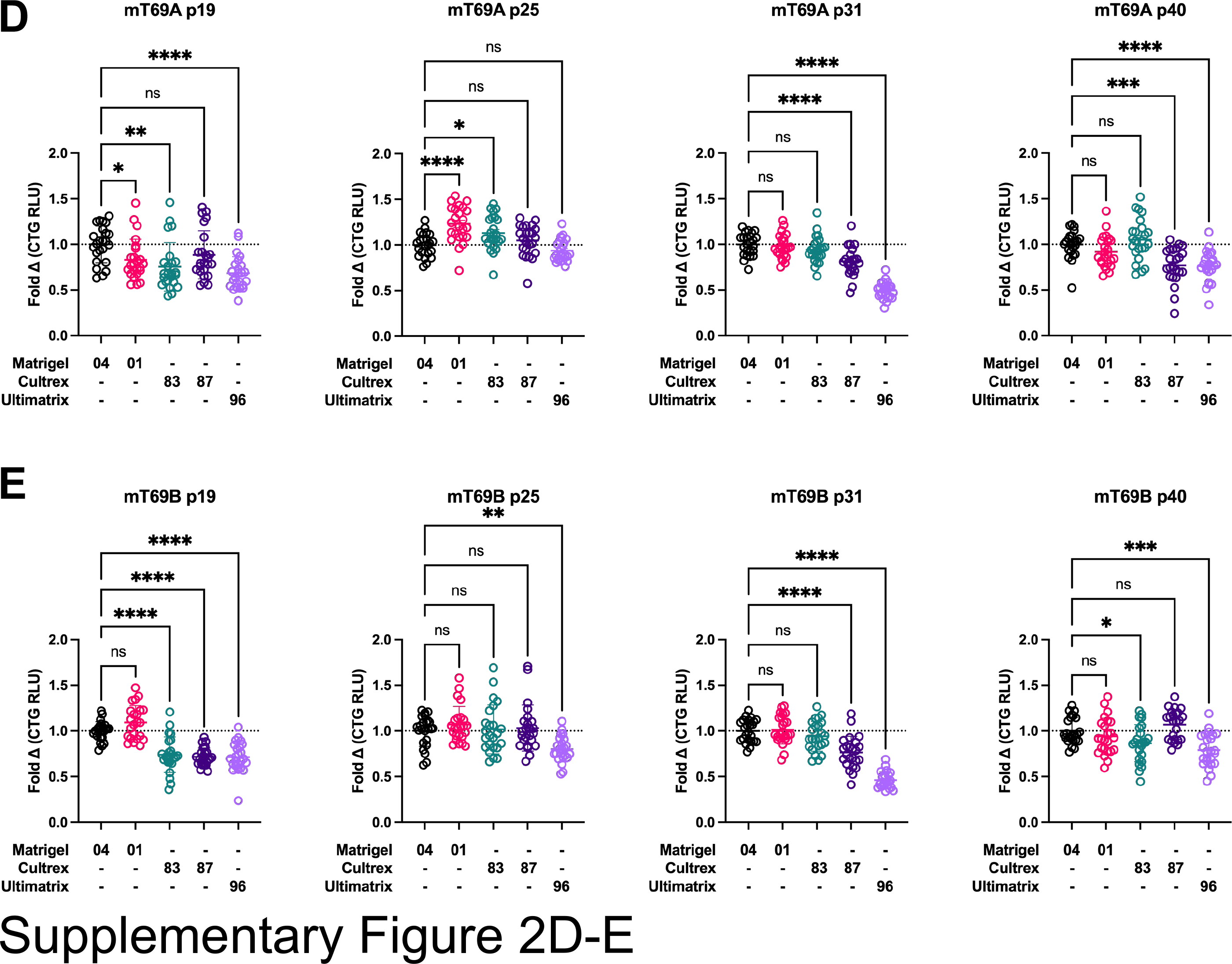
Proliferation of human and mouse organoids is not significantly altered by basement membrane extract. (**A**) Data for n = 3 experimental replicates of hF24 proliferation in Matrigel 04, Matrigel 01, Cultrex 83, Cultrex 87, or UltiMatrix 96. Data represent mean ± SD. \*\**P* ≤ 0.01, \*\*\*\**P* ≤ 0.0001 vs. Matrigel 04. (**B**) Data for n = 3 experimental replicates of hM1F proliferation in Matrigel 04, Matrigel 01, Cultrex 83, Cultrex 87, or UltiMatrix 96. Data represent mean ± SD. \*\**P* ≤ 0.01, \*\*\*\**P* ≤ 0.0001 vs. Matrigel 04. (**C**) Data for n = 2 experimental replicates of hT1 proliferation in Matrigel 04, Matrigel 01, Cultrex 83, Cultrex 87, or UltiMatrix 96. Data represent mean ± SD. \*\*\*\**P* ≤ 0.0001 vs. Matrigel 04. (**D**) Data for n = 4 experimental replicates of mT69A proliferation in Matrigel 04, Matrigel 01, Cultrex 83, Cultrex 87, or UltiMatrix 96. Data represent mean ± SD. \**P* ≤ 0.05, \*\**P* ≤ 0.01, \*\*\*\**P* ≤ 0.0001 vs. Matrigel 04. (**E**) Data for n = 4 experimental replicates of mT69B proliferation in Matrigel 04, Matrigel 01, Cultrex 83, Cultrex 87, or UltiMatrix 96. Data represent mean ± SD. \*\**P* ≤ 0.01, \*\*\*\**P* ≤ 0.0001 vs. Matrigel 04.

**Supplementary Figure 3.**
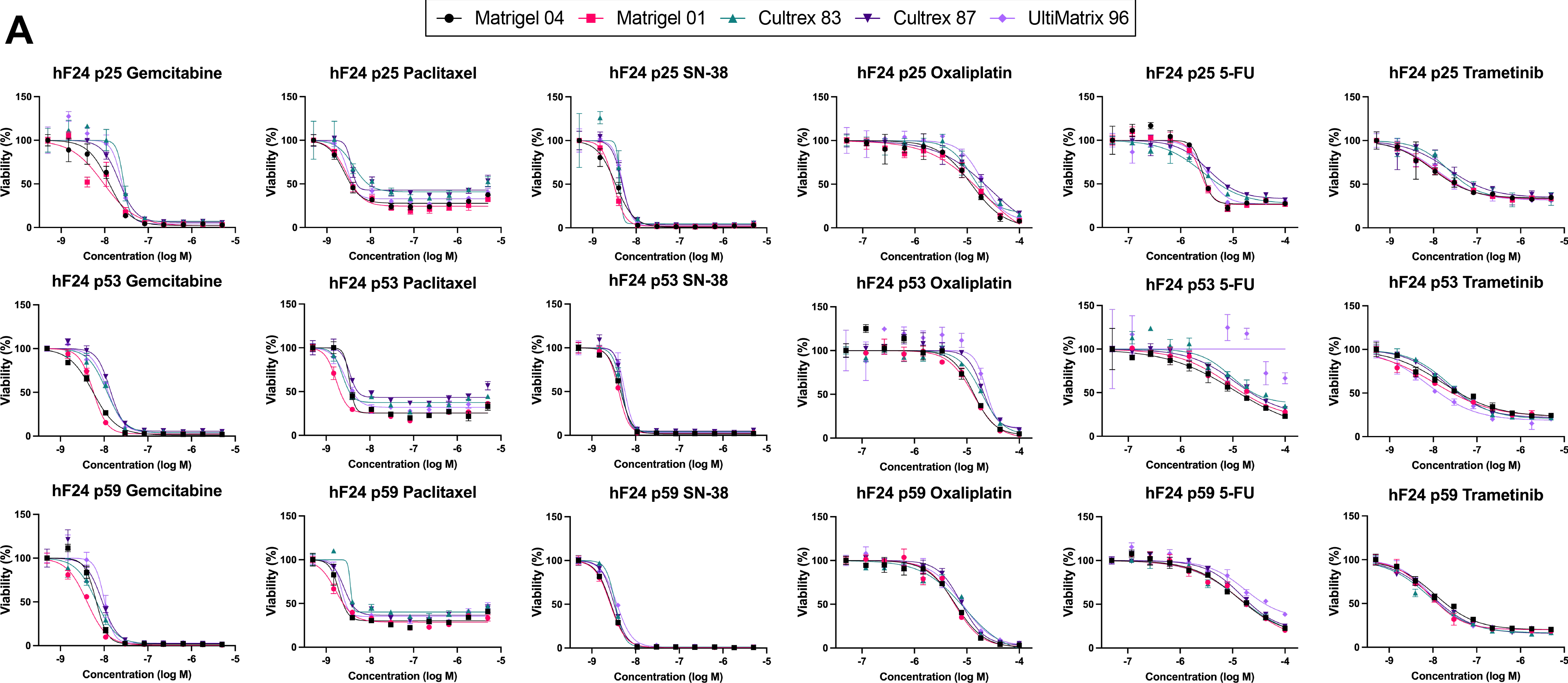
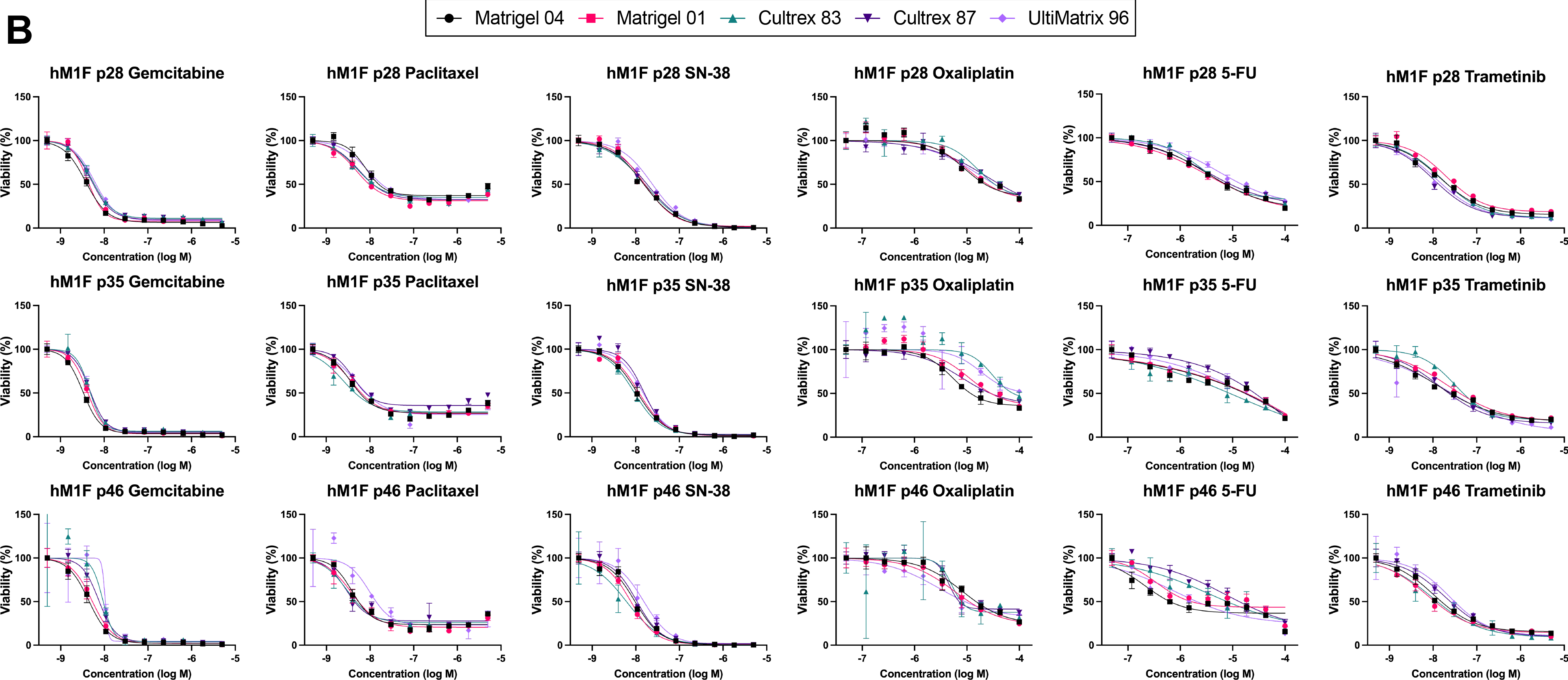
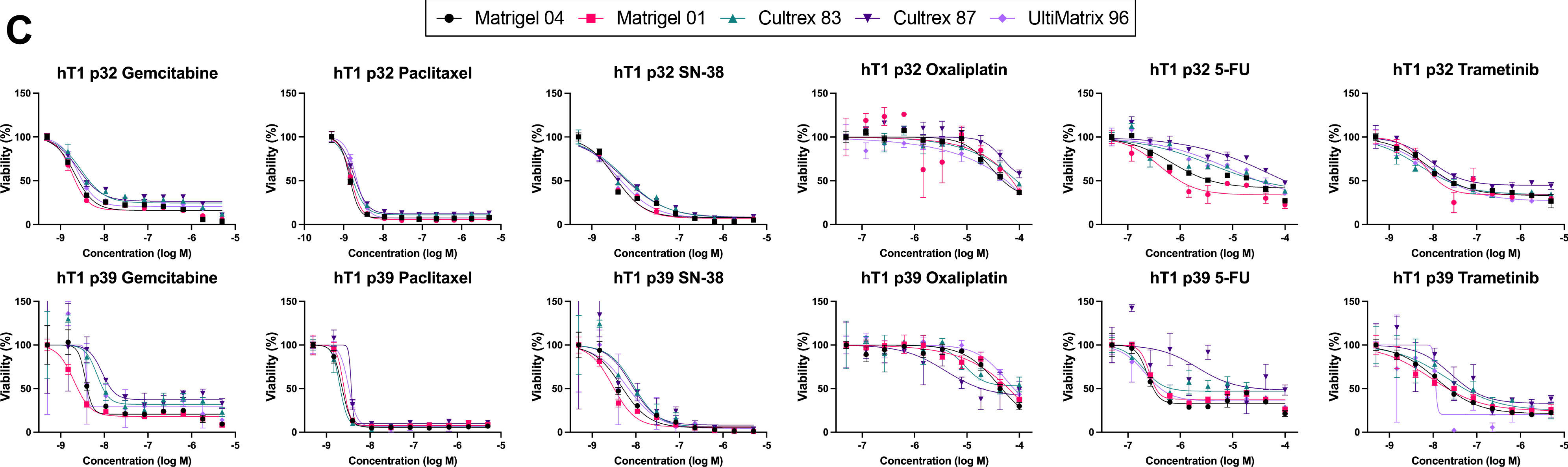
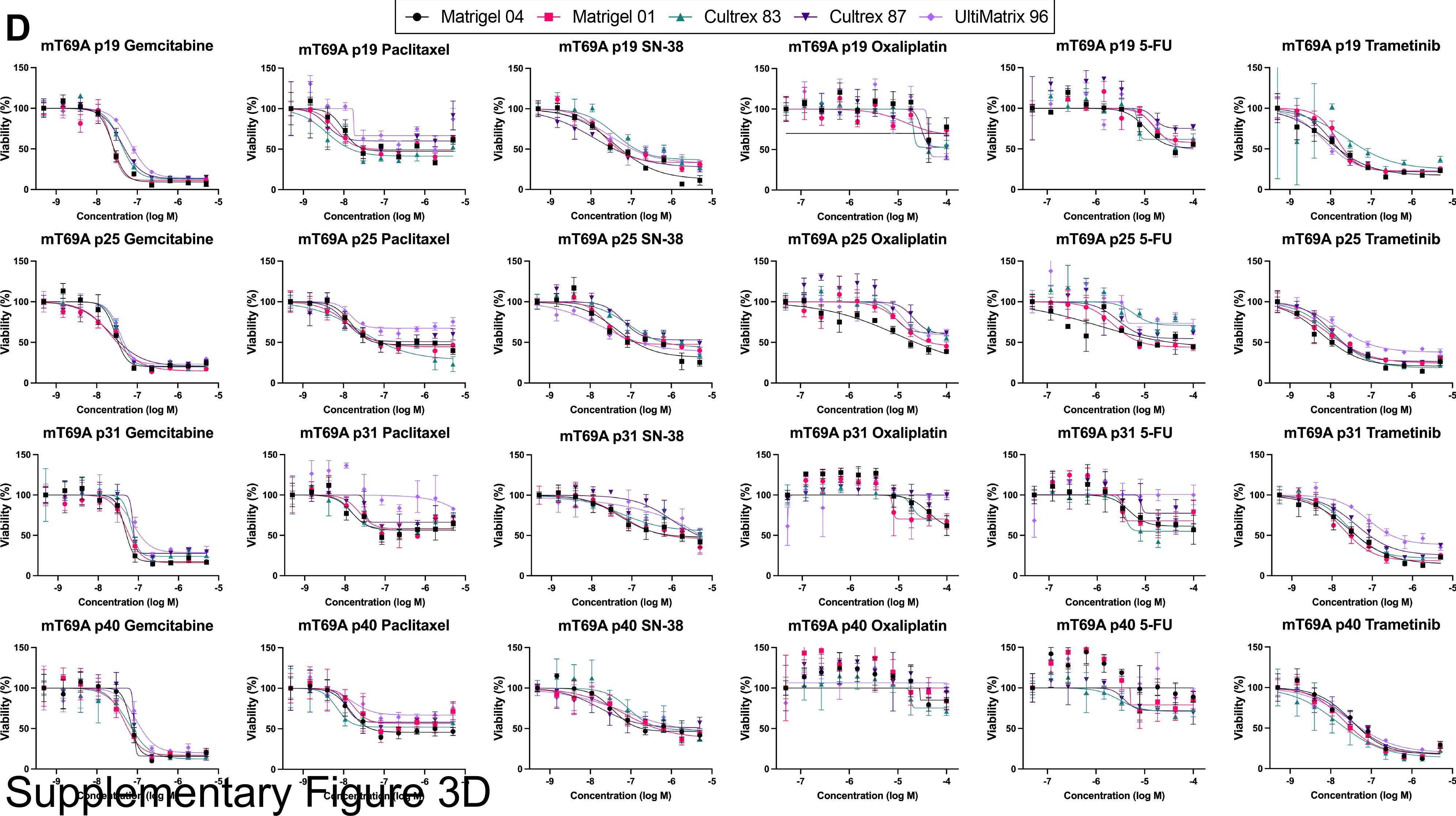
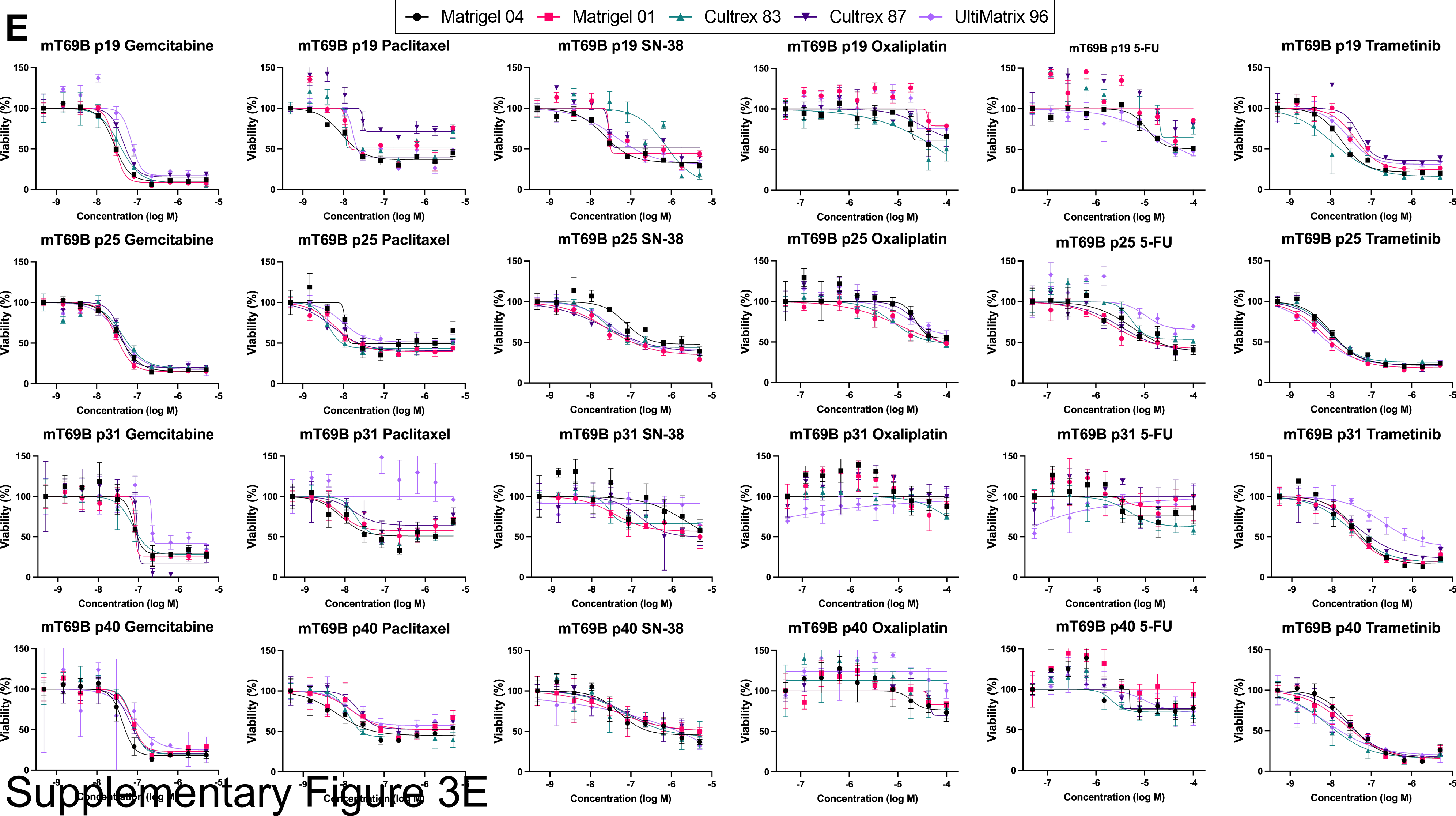
Dose response curves are consistent regardless of basement membrane extract used. (**A**) Data for n = 3 experimental replicates of hF24 response to gemcitabine, paclitaxel, SN-38, oxaliplatin, 5-FU, and trametinib while cultured in Matrigel 04, Matrigel 01, Cultrex 83, Cultrex 87, or UltiMatrix 96. Data represent mean ± SD. (**B**) Data for n = 3 experimental replicates of hM1F response to gemcitabine, paclitaxel, SN-38, oxaliplatin, 5-FU, and trametinib while cultured in Matrigel 04, Matrigel 01, Cultrex 83, Cultrex 87, or UltiMatrix 96. Data represent mean ± SD. (**C**) Data for n = 2 experimental replicates of hT1 response to gemcitabine, paclitaxel, SN-38, oxaliplatin, 5-FU, and trametinib while cultured in Matrigel 04, Matrigel 01, Cultrex 83, Cultrex 87, or UltiMatrix 96. Data represent mean ± SD. (**D**) Data for n = 4 experimental replicates of mT69A response to gemcitabine, paclitaxel, SN-38, oxaliplatin, 5-FU, and trametinib while cultured in Matrigel 04, Matrigel 01, Cultrex 83, Cultrex 87, or UltiMatrix 96. Data represent mean ± SD. (**E**) Data for n = 4 experimental replicates of mT69B response to gemcitabine, paclitaxel, SN-38, oxaliplatin, 5-FU, and trametinib while cultured in Matrigel 04, Matrigel 01, Cultrex 83, Cultrex 87, or UltiMatrix 96. Data represent mean ± SD.

**Supplementary Figure 4.**
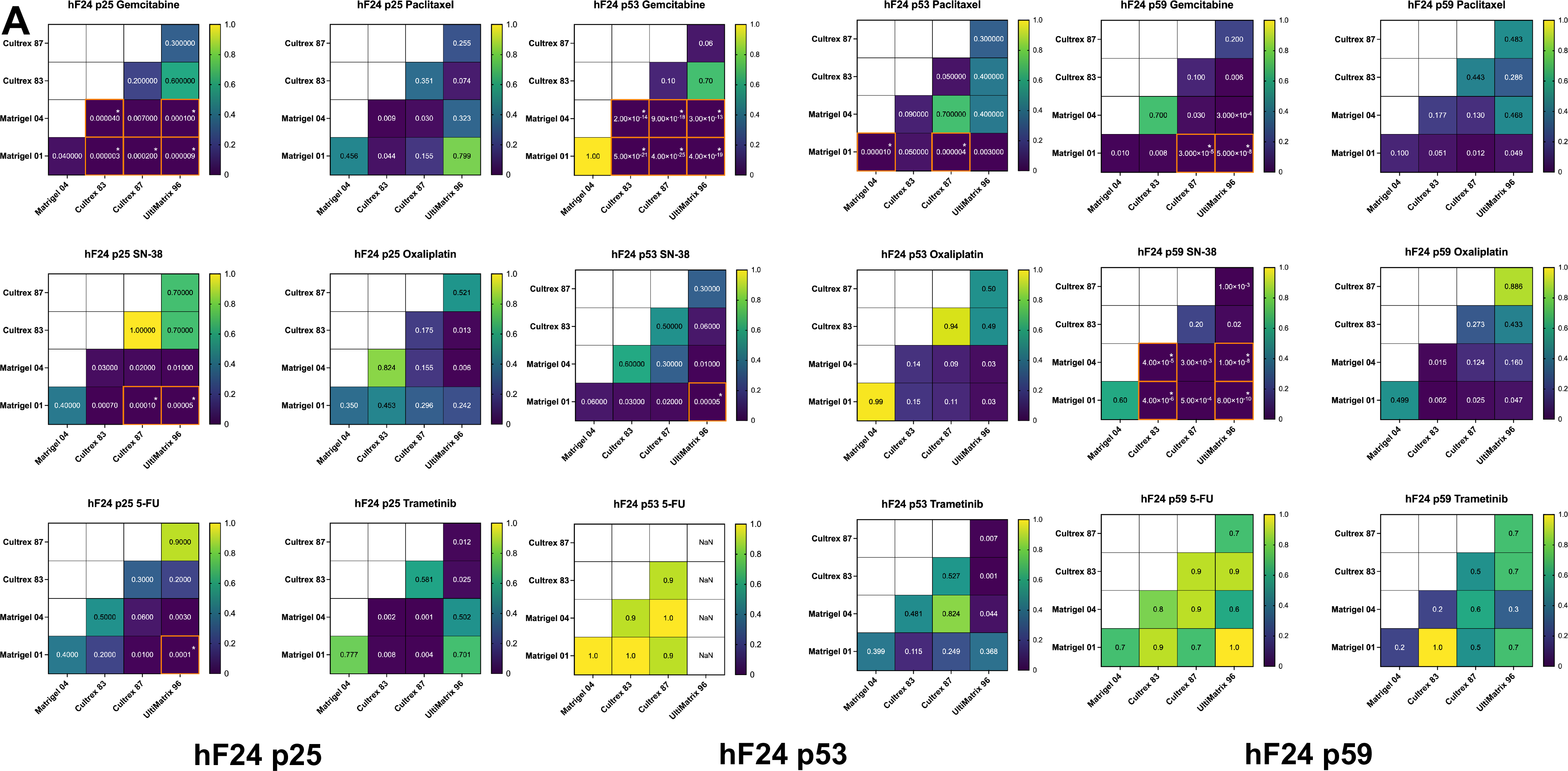
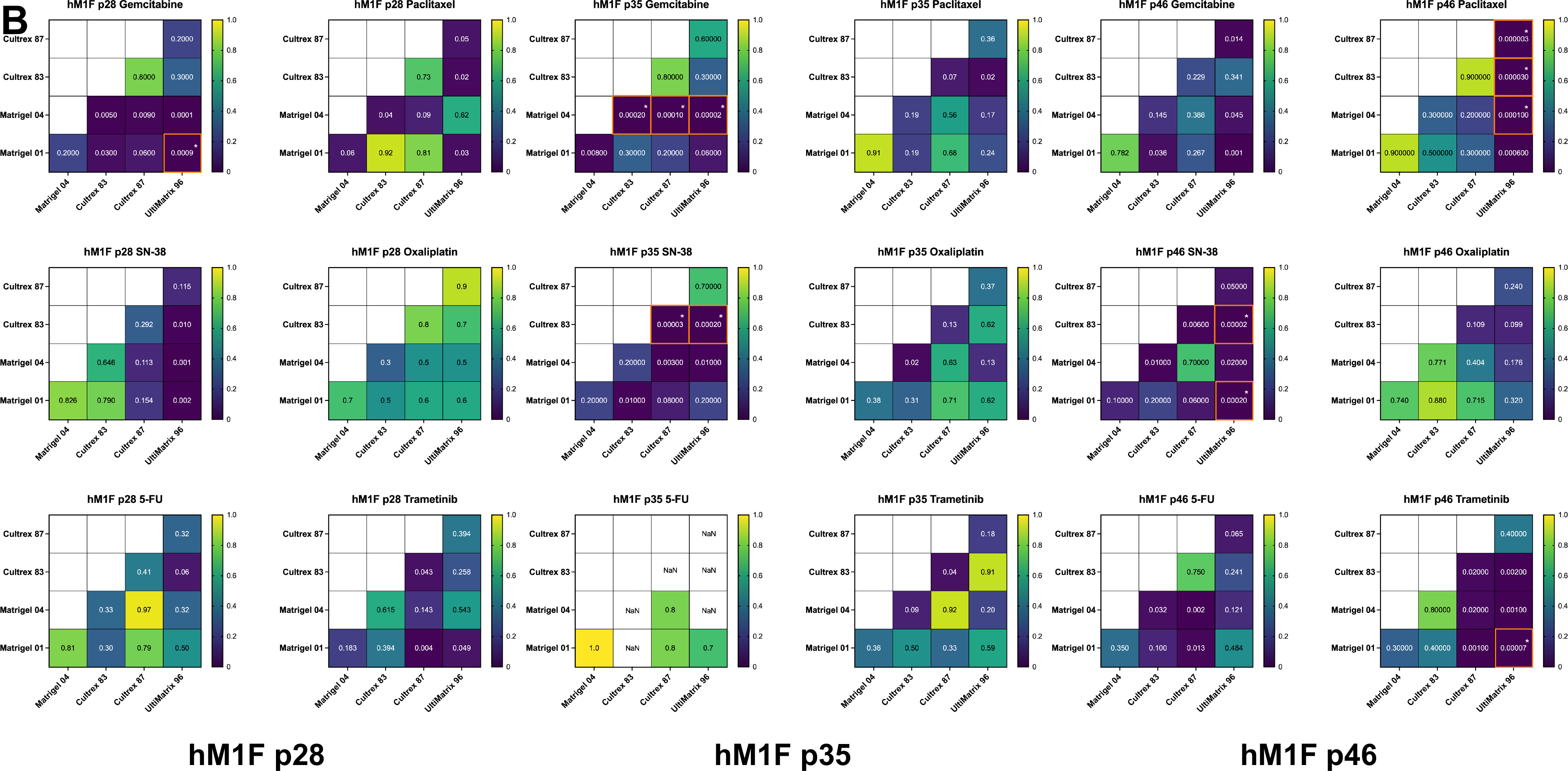
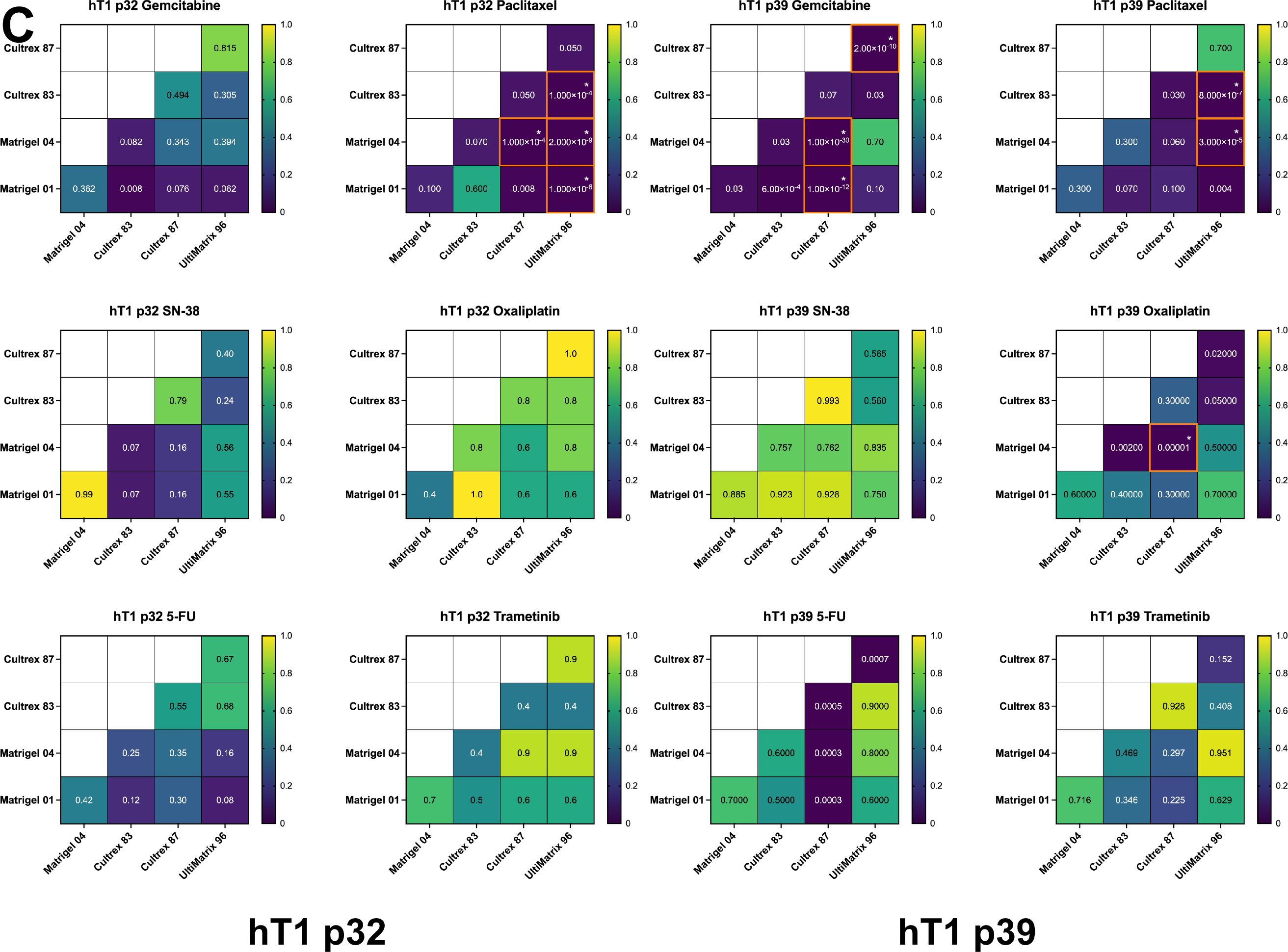
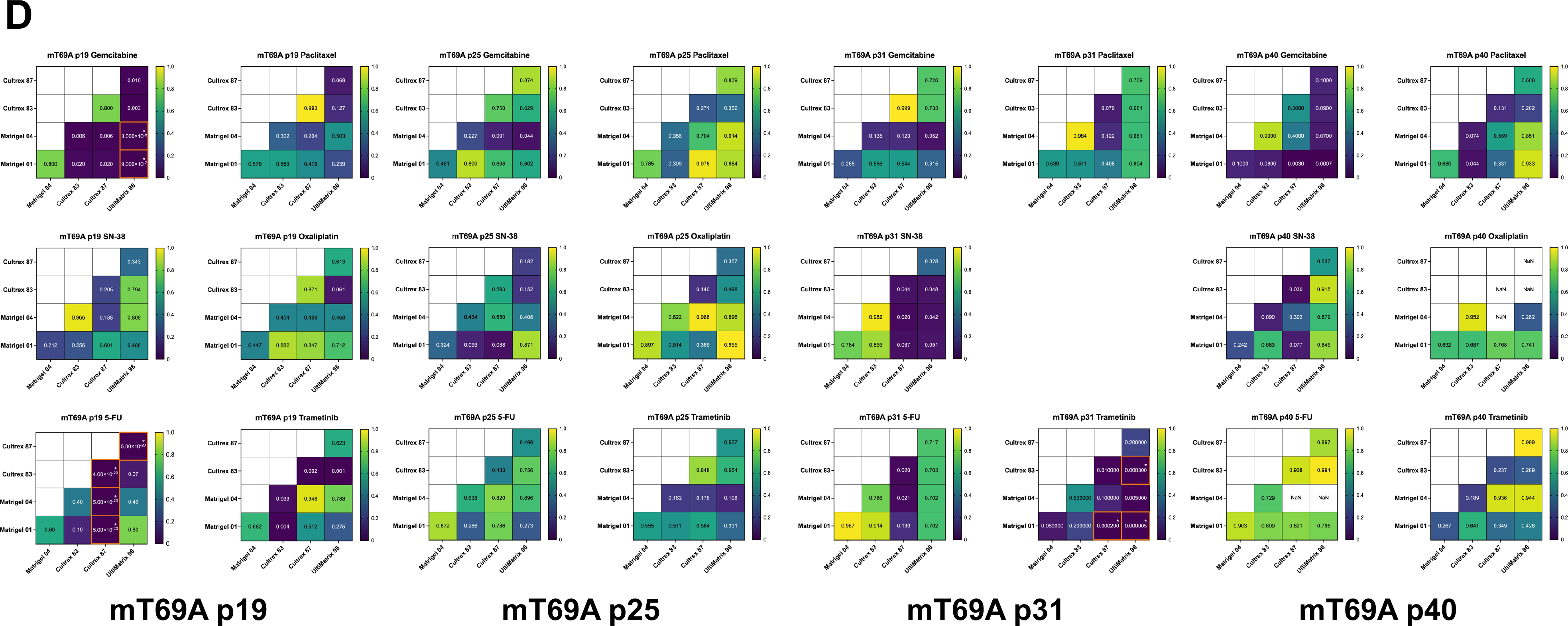
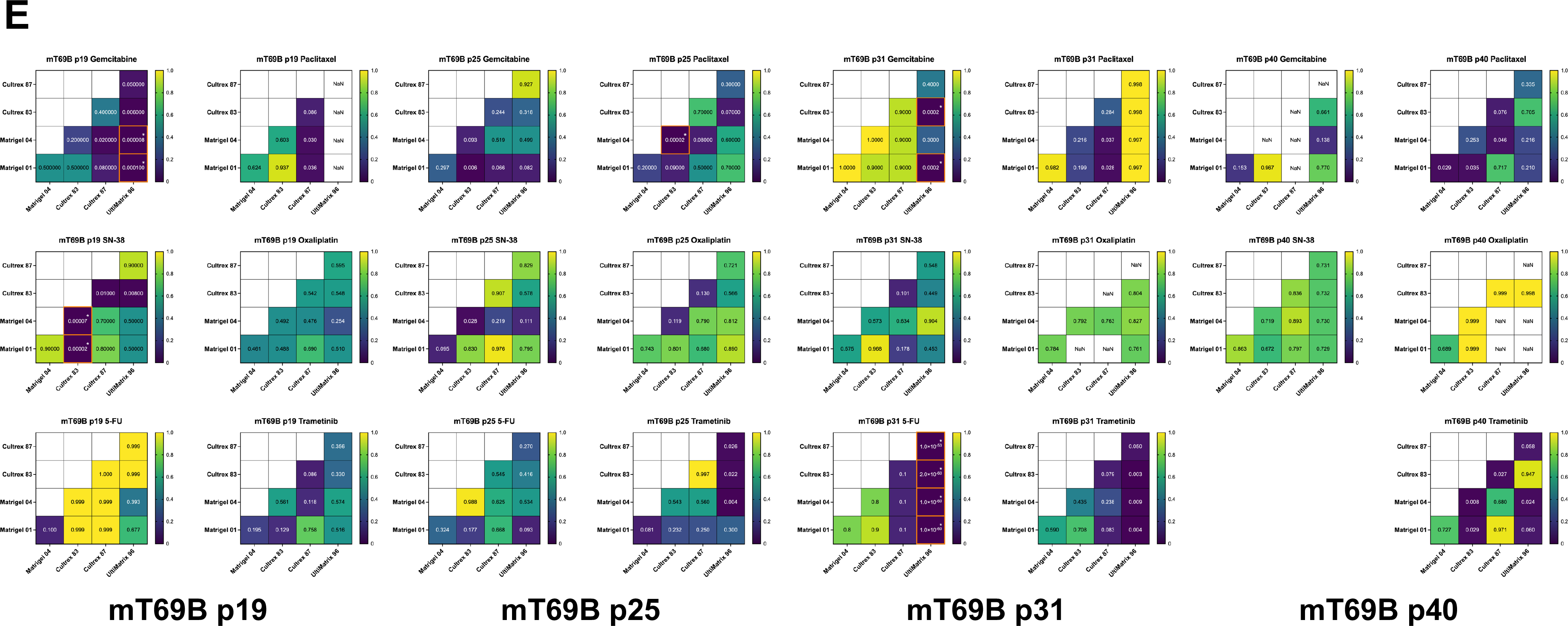
Statistically significant differences in IC50 within experimental replicates occur on a per-drug and per-organoid basis. (**A-C**) Heatmaps of *P* values from pairwise comparisons of IC50 between BMEs for each experimental repeat of hF24 (**A**), hM1F (**B**), or hT1 (**C**). Each square represents significance of pairwise t-tests. After Bonferroni adjustment, t-tests were considered significant if *P* < 0.00027778. Statistically significant comparisons are denoted by an asterisk and orange border. (**D-E**) Heatmaps of *P* values from pairwise comparisons of IC50 between BMEs for each experimental repeat of mT69A (**D**) or mT69B (**E**). Each square represents significance of pairwise t-tests. After Bonferroni adjustment, t-tests were considered significant if *P* < 0.00041667. Statistically significant comparisons are denoted by an asterisk and orange border.

**Supplementary Figure 5.**
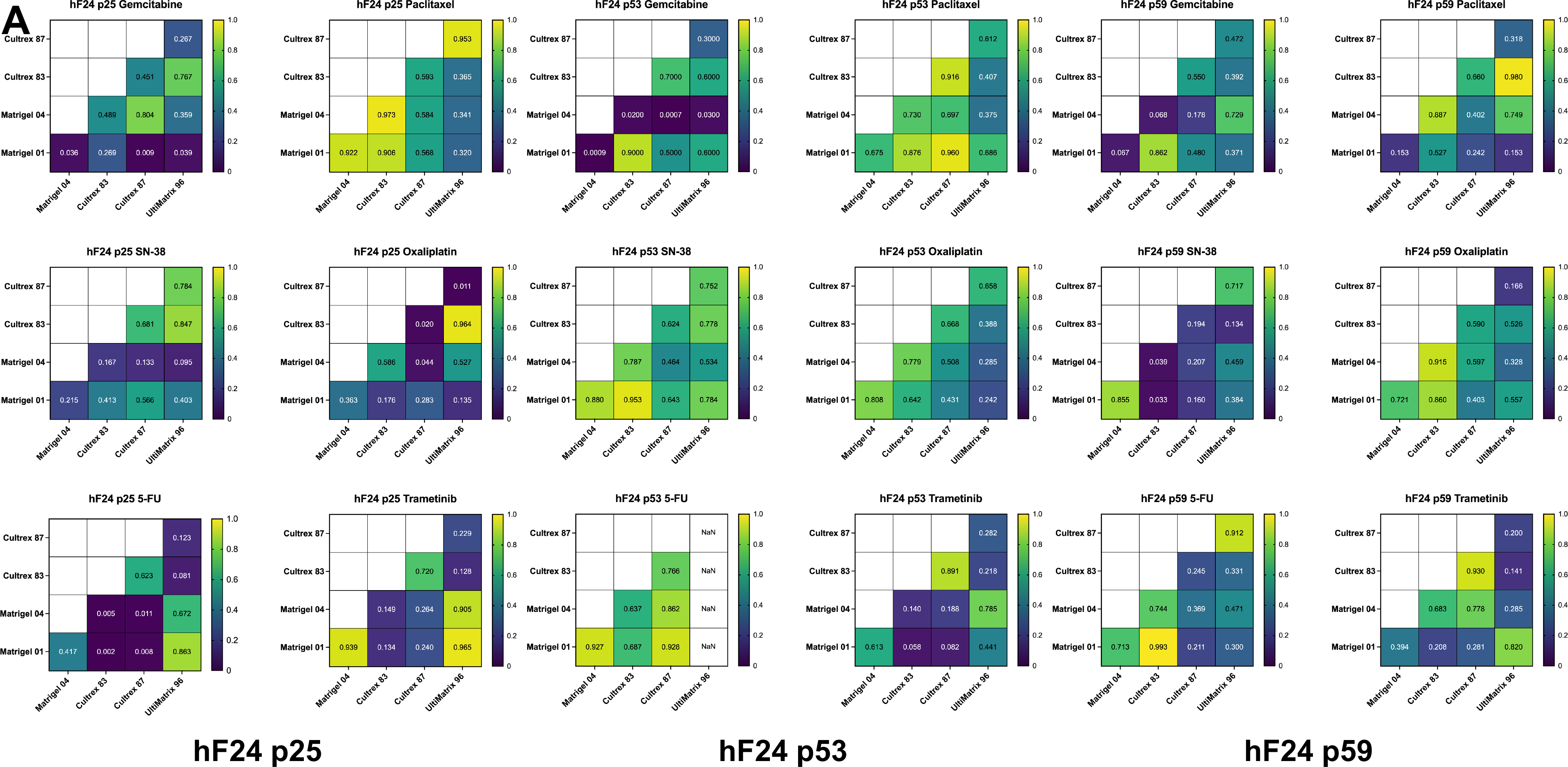
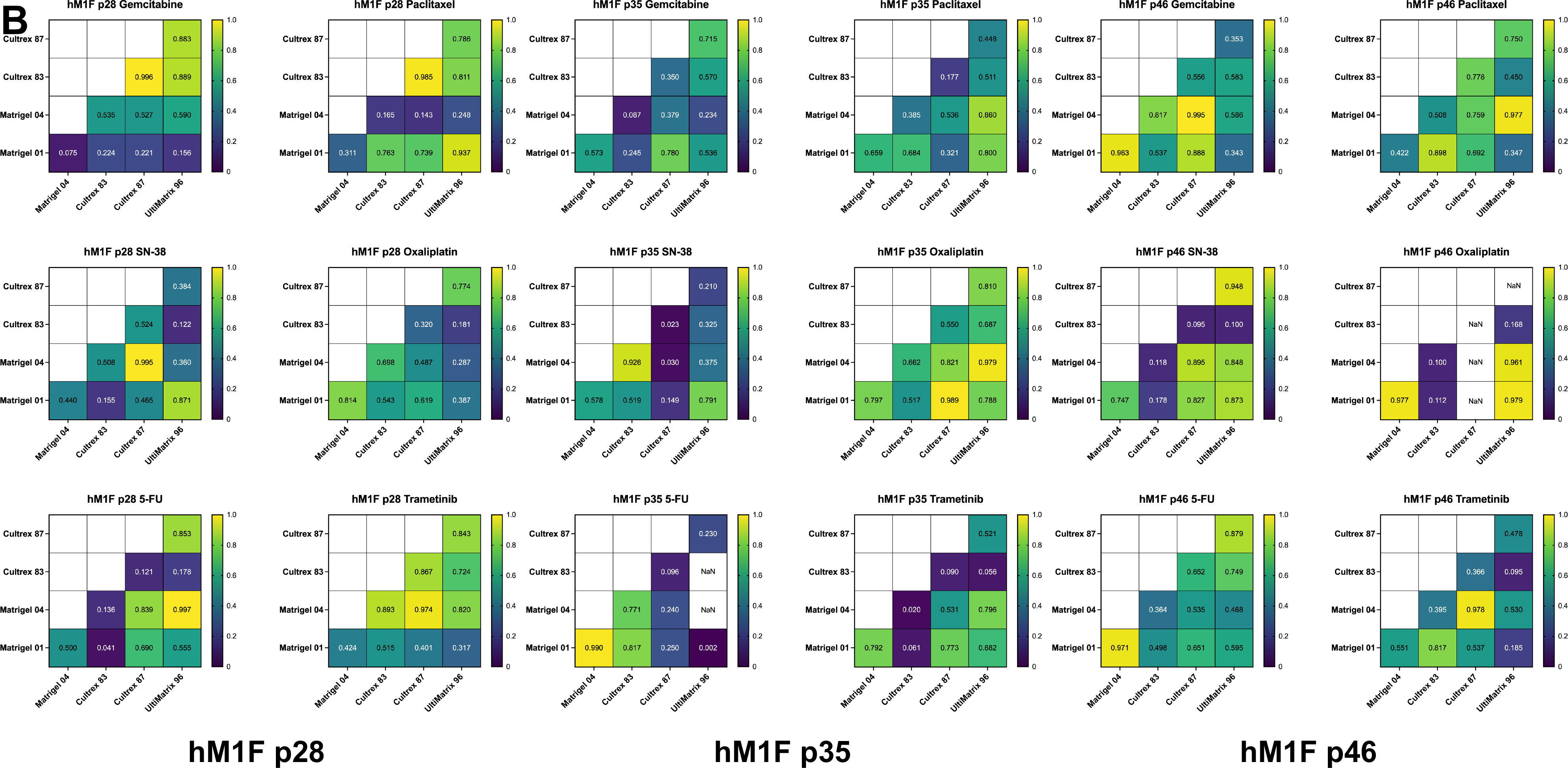
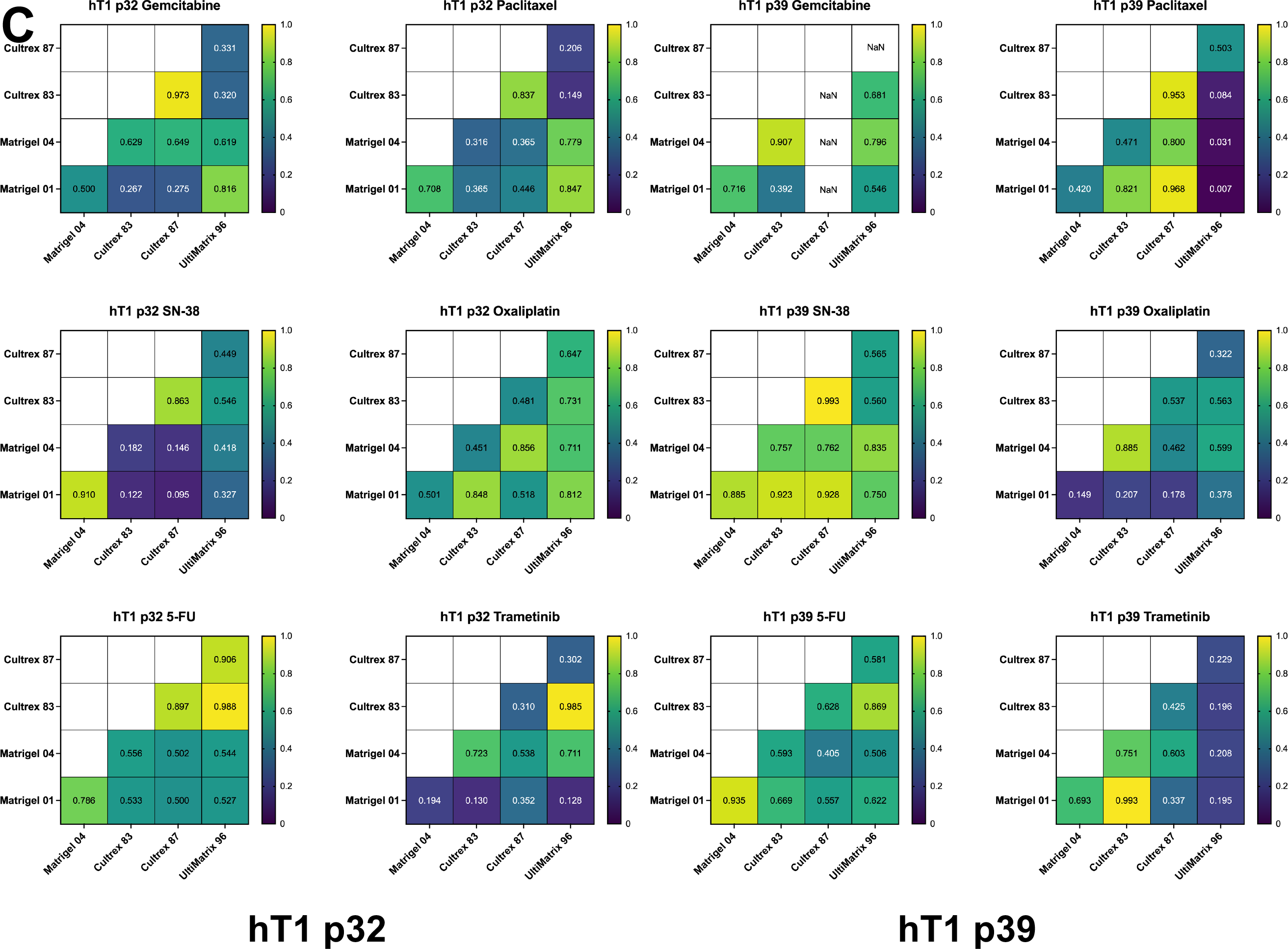
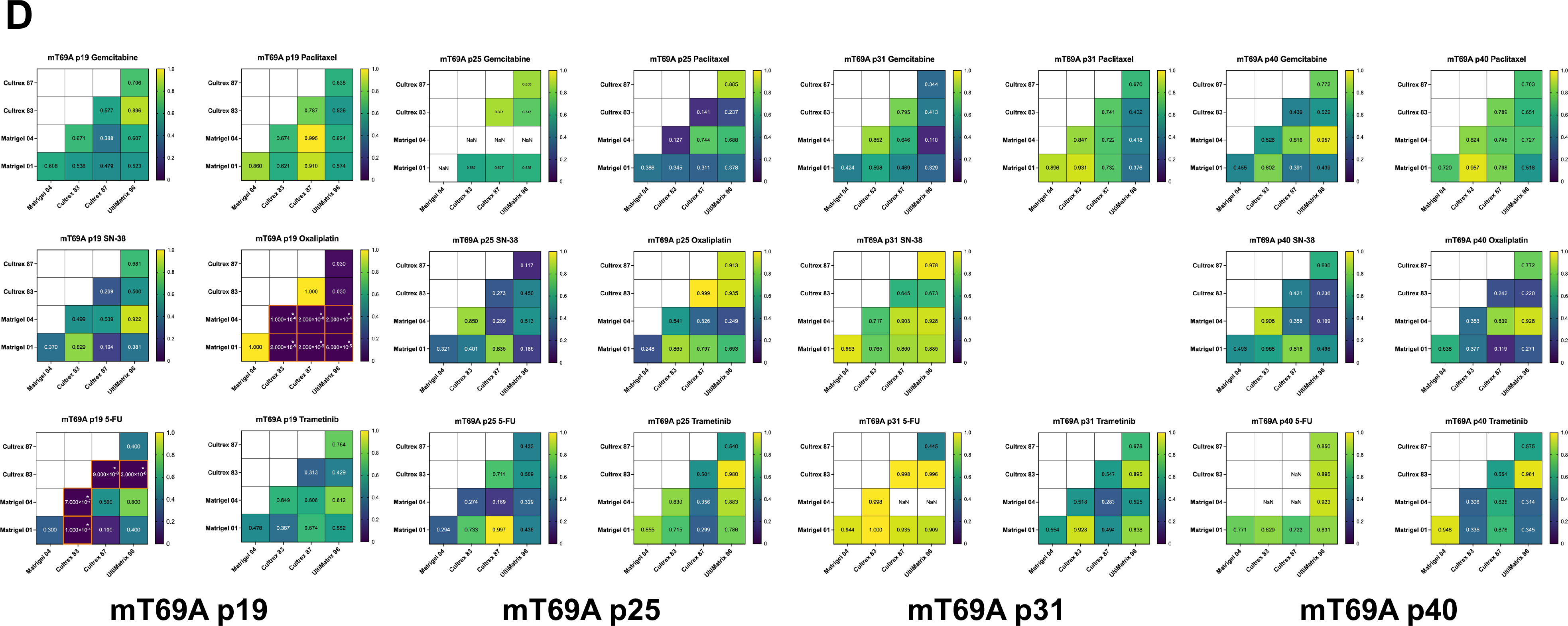
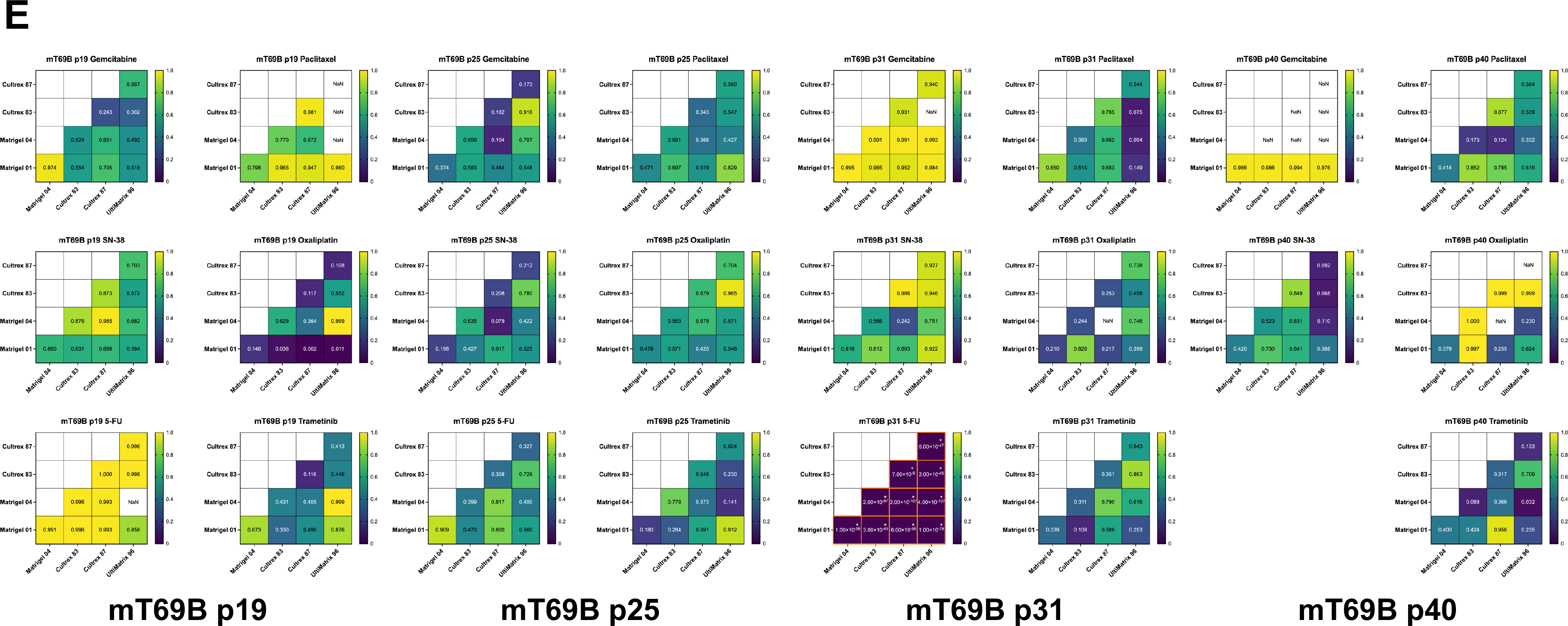
Dose response curve Hill slopes remain consistent across basement membrane extract culture conditions. (**A-C**) Heatmaps of *P* values from pairwise comparisons of Hill slope between BMEs for each experimental repeat of hF24 (**A**), hM1F (**B**), or hT1 (**C**). Each square represents significance of pairwise t-tests. After Bonferroni adjustment, t-tests were considered significant if *P* < 0.00027778. Statistically significant comparisons are denoted by an asterisk and orange border. (**D-E**) Heatmaps of *P* values from pairwise comparisons of Hill slope between BMEs for each experimental repeat of mT69A (**D**) or mT69B (**E**). Each square represents significance of pairwise t-tests. After Bonferroni adjustment, t-tests were considered significant if *P* < 0.00041667. Statistically significant comparisons are denoted by an asterisk and orange border.

**Supplementary Figure 6.**
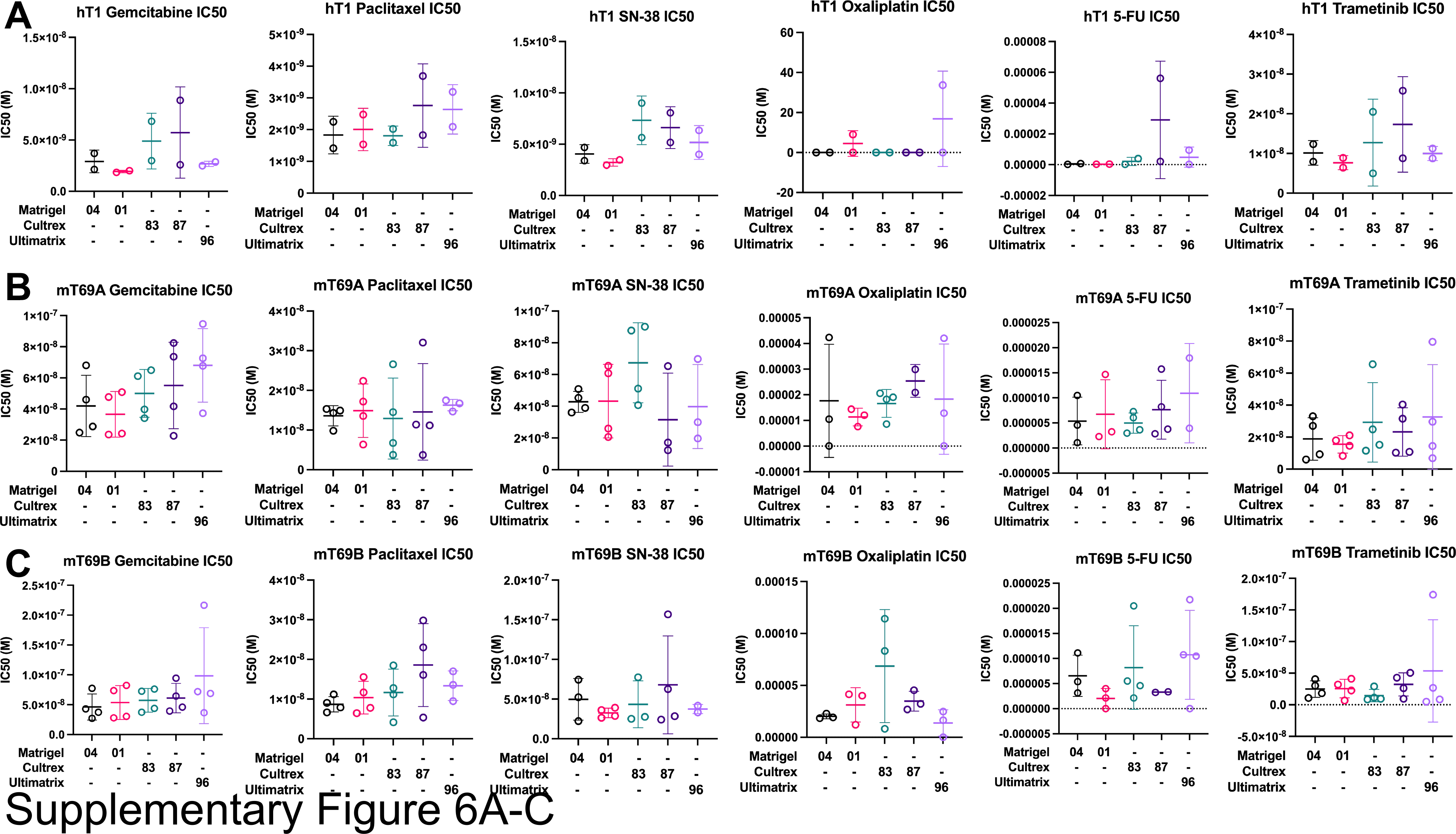
IC50 values remain consistent across basement membrane extract culture conditions. (**A**) IC50 values (M) for hT1 treated with gemcitabine, paclitaxel, SN- 38, oxaliplatin, 5-FU, and trametinib during culture in Matrigel 04, Matrigel 01, Cultrex 83, Cultrex 87, or UltiMatrix 96. Data represent mean ± SD of 2 experimental replicates. (**B**) IC50 values (M) for mT69A treated with gemcitabine, paclitaxel, SN-38, oxaliplatin, 5-FU, and trametinib during culture in Matrigel 04, Matrigel 01, Cultrex 83, Cultrex 87, or UltiMatrix 96. Data represent mean ± SD of 4 experimental replicates. (**C**) IC50 values (M) for mT69B treated with gemcitabine, paclitaxel, SN-38, oxaliplatin, 5-FU, and trametinib during culture in Matrigel 04, Matrigel 01, Cultrex 83, Cultrex 87, or UltiMatrix 96. Data represent mean ± SD of 4 experimental replicates. Outliers were determined using Grubbs’ test α = 0.05.

**Supplementary Figure 7.**
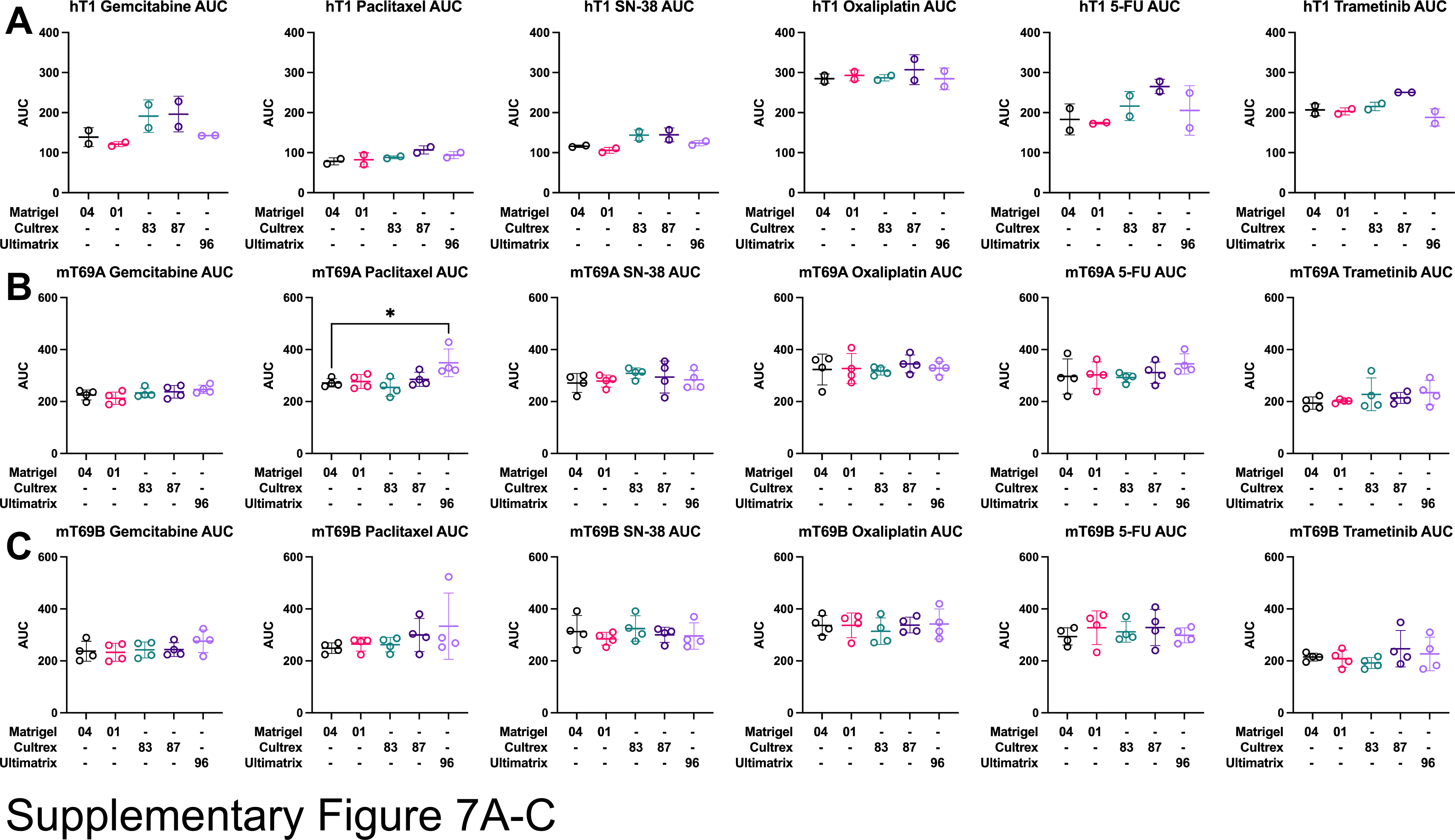
AUC values are consistent across basement membrane extract culture conditions. (**A**) AUC values for hT1 treated with gemcitabine, paclitaxel, SN-38, oxaliplatin, 5-FU, and trametinib during culture in Matrigel 04, Matrigel 01, Cultrex 83, Cultrex 87, or UltiMatrix 96. Data represent mean ± SD of 2 experimental replicates. (**B**) AUC values for mT69A treated with gemcitabine, paclitaxel, SN-38, oxaliplatin, 5-FU, and trametinib during culture in Matrigel 04, Matrigel 01, Cultrex 83, Cultrex 87, or UltiMatrix 96. Data represent mean ± SD of 4 experimental replicates. Statistical significance was determined by 1-way ANOVA. \**P* ≤ 0.05 vs. Matrigel 04. (**C**) AUC values for mT69B treated with gemcitabine, paclitaxel, SN-38, oxaliplatin, 5-FU, and trametinib during culture in Matrigel 04, Matrigel 01, Cultrex 83, Cultrex 87, or UltiMatrix 96. Data represent mean ± SD of 4 experimental replicates.

**Supplementary Figure 8.**
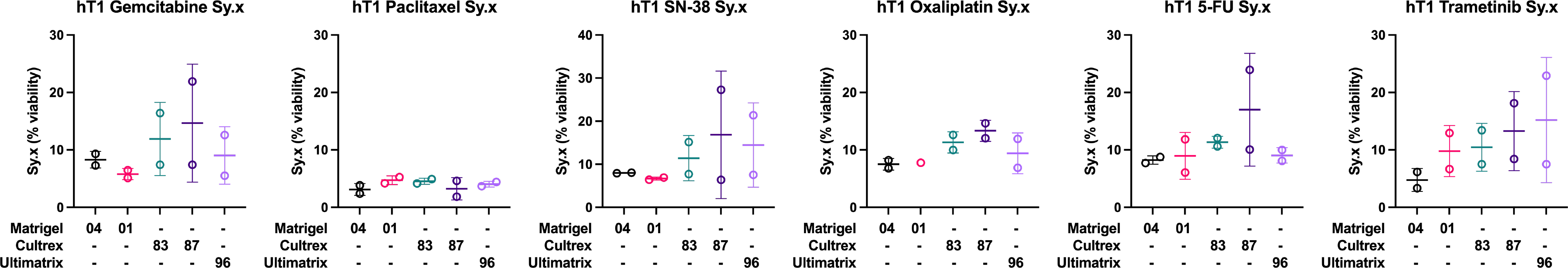
Standard deviation of the residuals does not statistically change as a function of basement membrane extract used. Standard deviation of the residuals (Sy.x) for hT1 treated with gemcitabine, paclitaxel, SN-38, oxaliplatin, 5-fluorourical (5-FU), and trametinib during culture in Matrigel 04, Matrigel 01, Cultrex 83, Cultrex 87, or UltiMatrix 96. Data represent mean ± SD of 2 experimental replicates. Outliers were determined using Grubbs’ test α = 0.05.

**Supplementary Figure 9.**
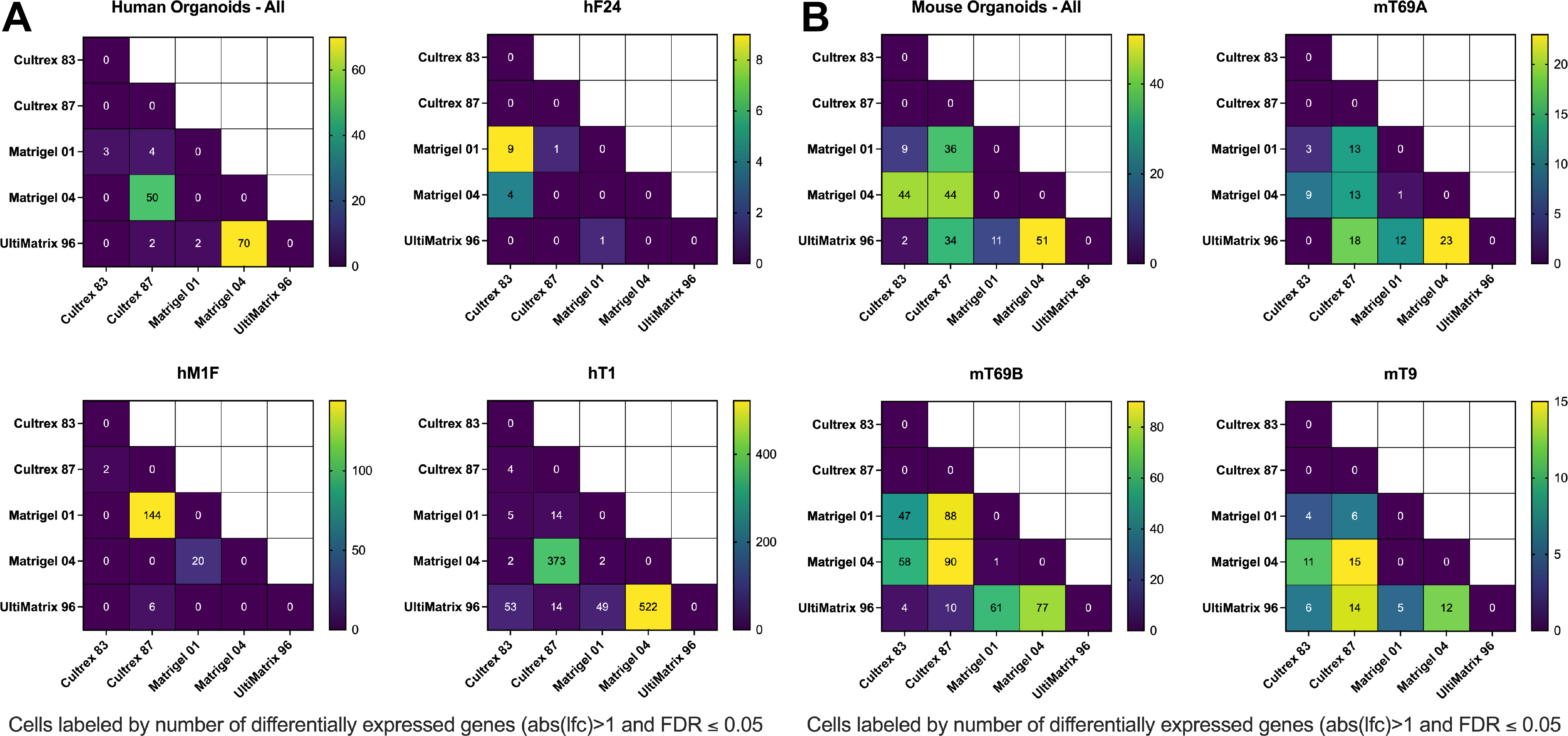
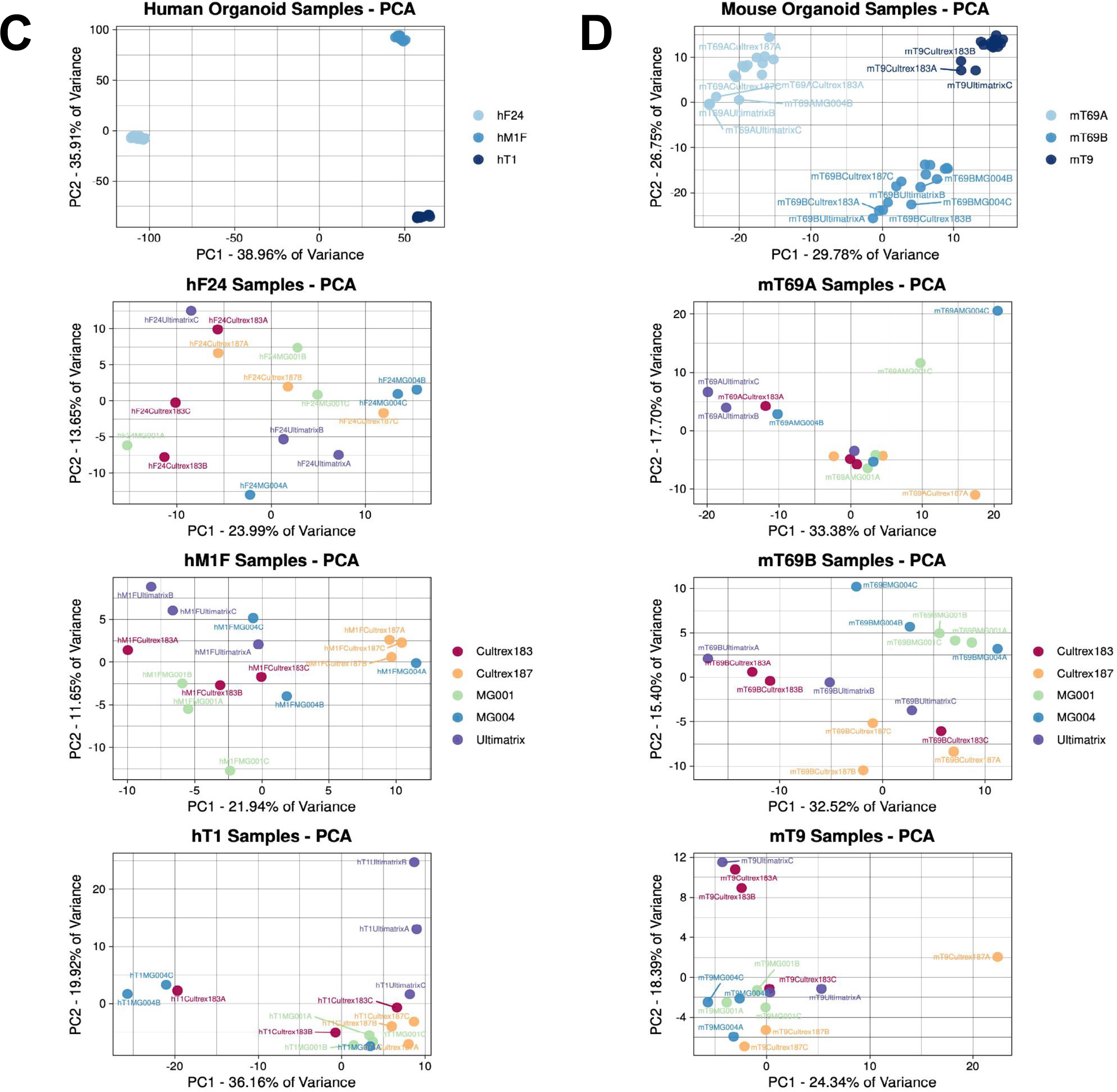
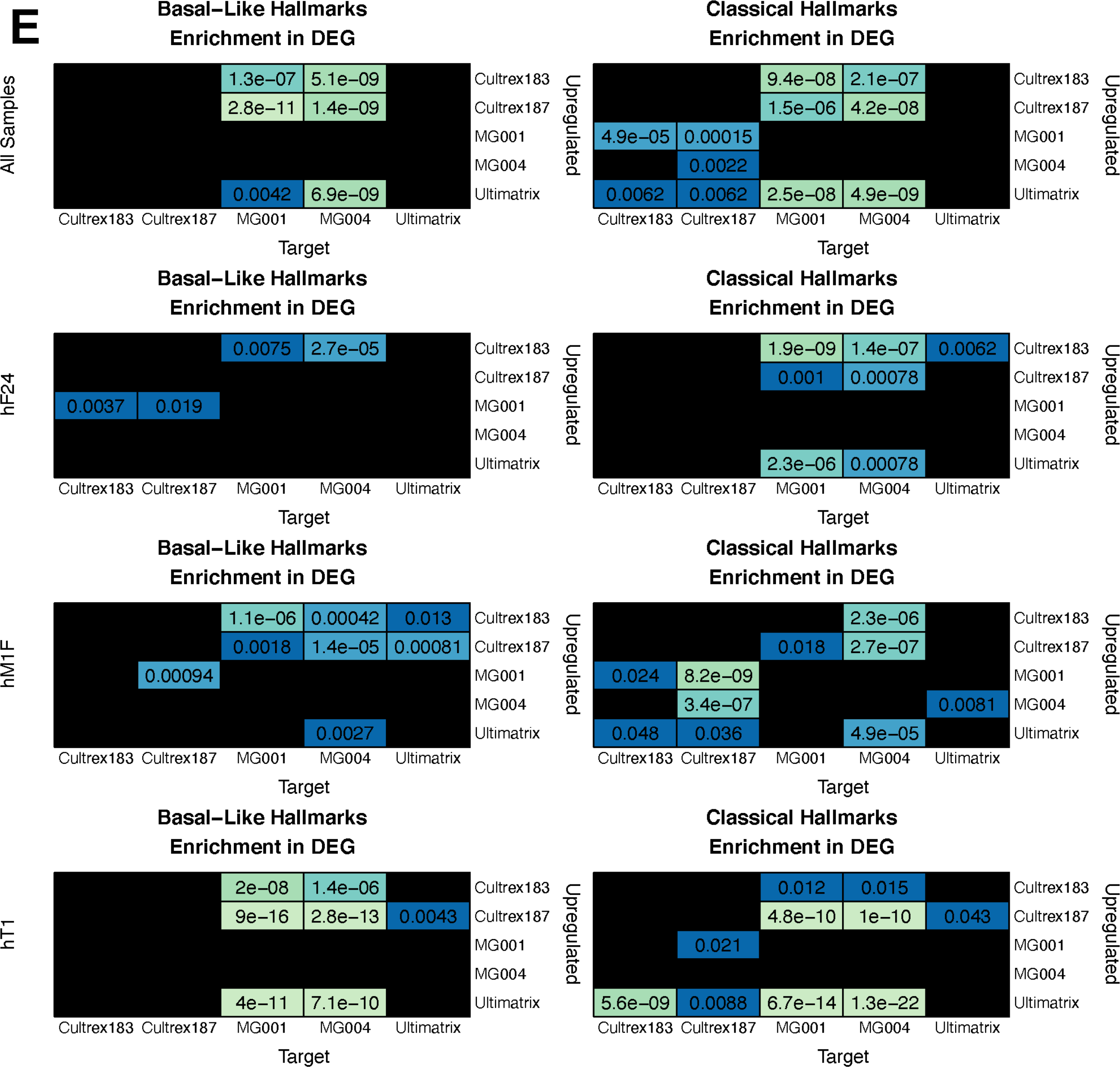
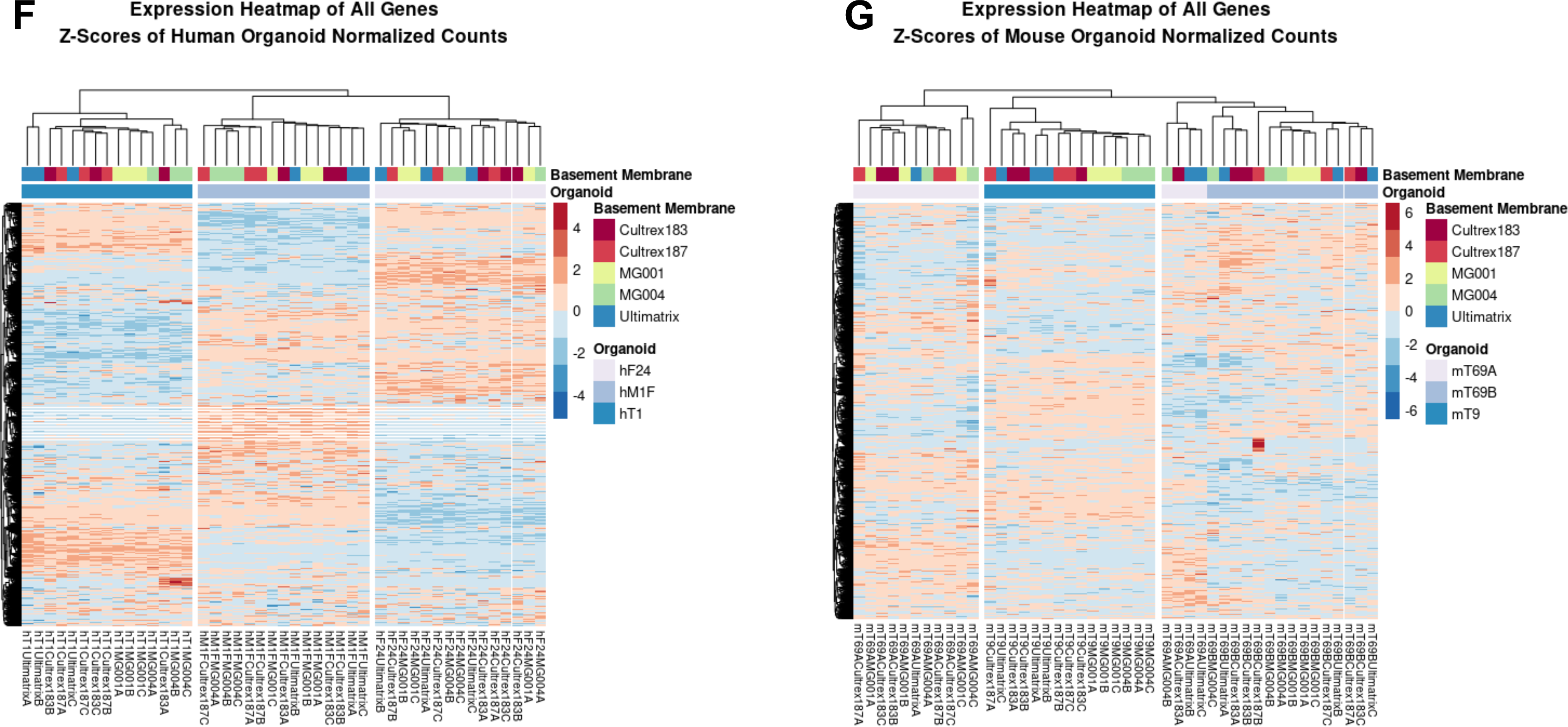
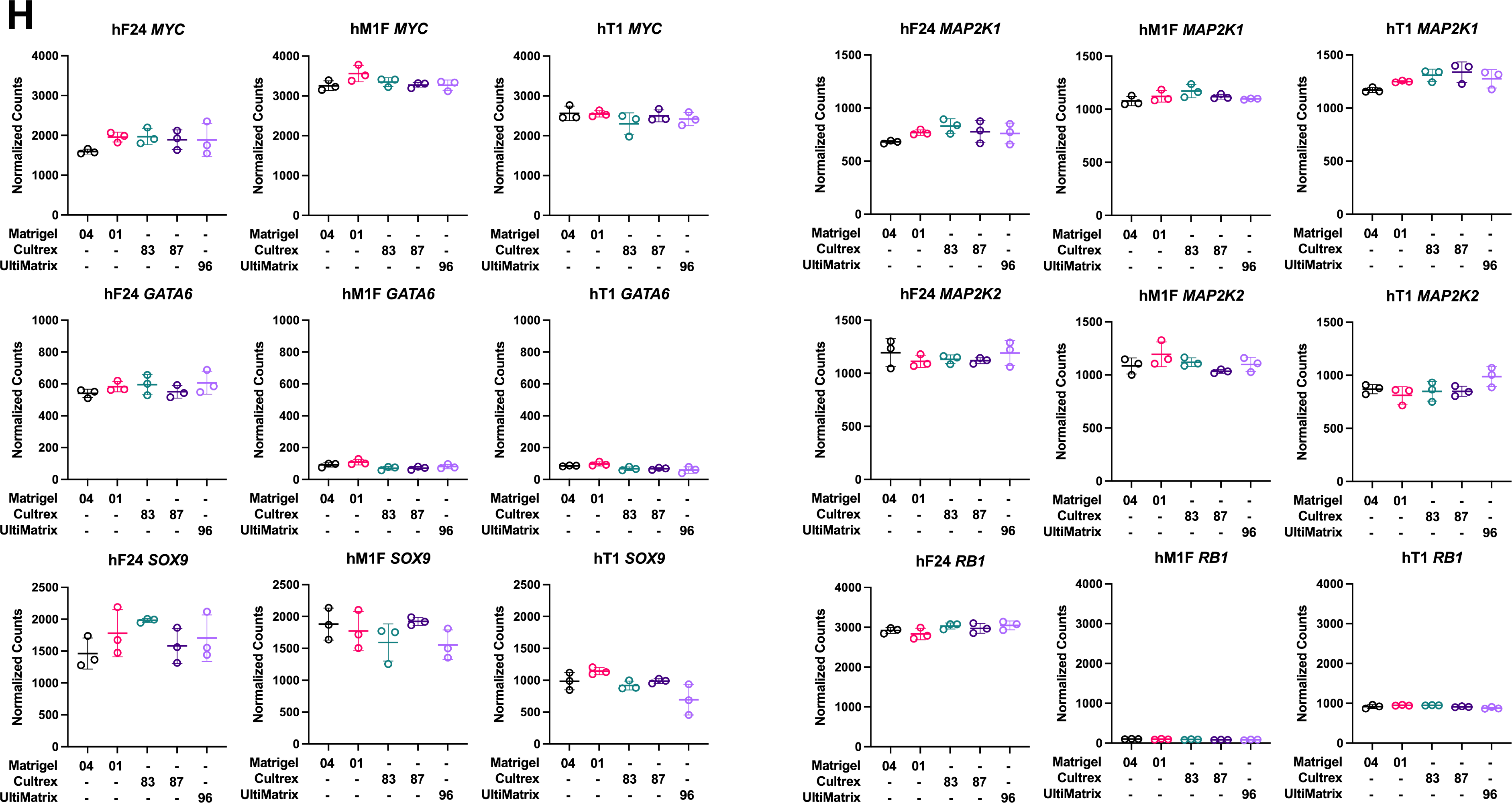
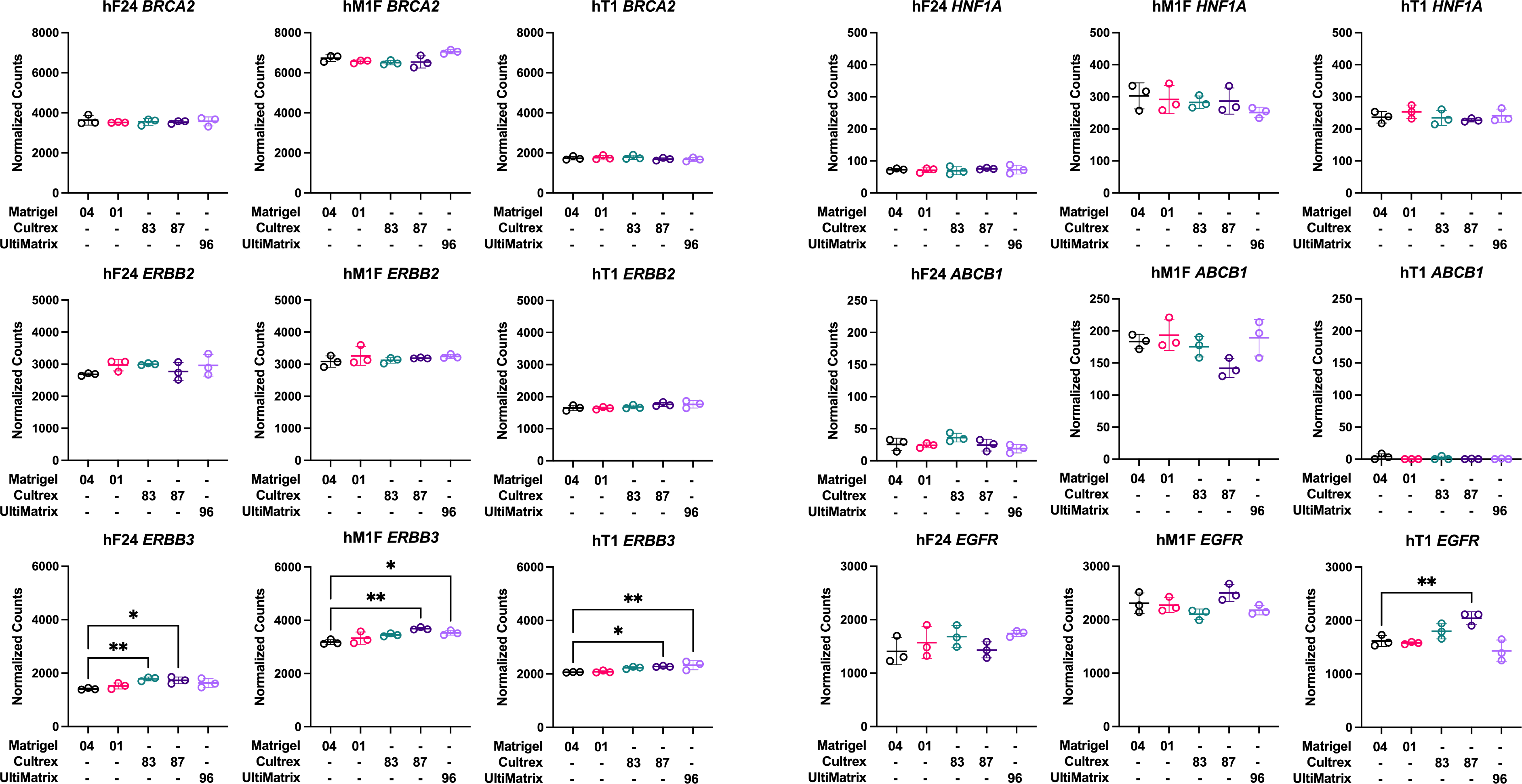
Global gene expression profiles are consistent across basement membrane extract conditions for both mouse and human organoids. (**A**) Differential gene expression among human patient-derived organoids cultured in Matrigel 04 (MG004), Matrigel 01 (MG001), Cultrex 83 (Cultrex 183), Cultrex 87 (Cultrex 187), or UltiMatrix 96 (Ultimatrix). Numbers represent total number of genes with an adjusted *P* value of differential expression ≤ 0.05 and an absolute log2(foldchange) > 1 for all pairwise comparisons of basement membrane extracts within individual organoid lines and within species. (**B**) Differential gene expression among mouse organoids cultured in Matrigel 04 (MG004), Matrigel 01 (MG001), Cultrex 83 (Cultrex 183), Cultrex 87 (Cultrex 187), or UltiMatrix 96 (Ultimatrix). Numbers represent total number of genes with an adjusted *P* value of differential expression ≤ 0.05 and an absolute log2(foldchange) > 1 for all pairwise comparisons of basement membrane extracts within individual organoid lines and within species. (**C**) Principal component analysis of human patient-derived organoid gene expression within species and individual organoid lines. (**D**) Principal component analysis of mouse organoid gene expression within species and individual organoid lines. (**E**) Heatmaps of basal-like or classical hallmark gene enrichment between basement membrane extract comparisons for all human patient-derived organoid lines (top row) and individual lines. Each square represents significance of enrichment in the differentially expressed genes between two basement membrane extracts. The differentially expressed genes in the BME listed on the y-axis are queried for enrichment. Squares are labeled with *P* value. Black squares = ns. (**F**) Expression heatmap of all genes in human patient-derived organoids cultured in different BMEs. Z-scores of normalized counts. (**G**) Expression heatmap of all genes in mouse organoids cultured in different basement membrane extracts. Z-scores of normalized counts. (**H**) Normalized counts for select genes of interest in hF24, hM1F, and hT1 cultured in different BMEs. Data represent mean ± SD of 3 technical replicates. \**P* ≤ 0.05, \*\**P* ≤ 0.01 vs. Matrigel 04.

Supplementary Table 1
| PDO | Origin | Site | Type of sample | Sex | Age | Race | Stage | KRAS (allele frequencies) | TP53 (allele frequencies) | SMAD4 (allele frequencies) | CDKN2A (allele frequencies) |
| --- | --- | --- | --- | --- | --- | --- | --- | --- | --- | --- | --- |
| hF24* | H. sapiens | Pancreas | EUS FNA | F | 57 | Caucasian | 4 | G12V 82% | F113V (100%) | LOSS (N/A) | DEEP LOSS (N/A) |
| hM1F* | H. sapiens | Lung | RAP | M | 55 | Caucasian | 4 | G12D 73% | R175H (100%) | WT (N/A) | WT (N/A) |
| hT1* | H. sapiens | Pancreas | Resection | M | 65 | Caucasian | 2B | G12V 39% | R213* (100%) | WT (N/A) | DEEP LOSS (N/A) |

\*H. Tiriac *et al.*, Organoid Profiling Identifies Common Responders to Chemotherapy in Pancreatic Cancer. *Cancer Discov* **8**, 1112-1129 (2018).

Supplementary Table 2
| PDO | Origin | Sex | Site | Type of sample |
| --- | --- | --- | --- | --- |
| mT69A | M. musculus | F | Pancreas | KPC B6, PDAC local pancreatic metastasis |
| mT69B | M. musculus | F | Pancreas | KPC B6, PDAC primary tumor |
| mT9 | M. musculus | F | Pancreas | KPC PDAC primary tumor |

## Notes

### Competing Interest Statement

The authors have declared no competing interest.

### Summary of Updates

Additional statistical analysis was conducted on dose-response curve parameters. The main text underwent revisions for clarity and wording. Figures were also revised to be color-blind safe, to include additional data from subsequent statistical analyses, and for clarity. A supplemental file including gene lists for classical- and basal-like-associated genes is also included.

